# Two Ways to Change Your Mind: Effects of Intentional Strength and Motor Costs on Changes of Intention

**DOI:** 10.1101/841882

**Authors:** Anne Löffler, Anastasia Sylaidi, Zafeirios Fountas, Patrick Haggard

**Author notes:** Corresponding Author: Anne Löffler, Zuckerman Mind Brain Behaviour Institute, Columbia University, 3227 Broadway, New York, NY 10027, USA.

## Abstract

Changes of Mind are a striking example of the human ability to flexibly reverse decisions after commitment to an initial choice, and to change actions according to circumstances. Previous studies of Changes of Mind largely focused on perceptual choices. Here we investigate reversals of *voluntary, endogenous* action decisions. In a novel version of the random-dot motion task, participants moved to a target that matched both the perceived dot-motion direction and an internally-generated intention (which colour target to move to). Movement trajectories revealed whether and when participants 1) perceived a change in dot-motion direction, or additionally 2) changed the colour that they chose to move to (‘Change of Intention’). Changes of Intention were less frequent in participants with strong colour intentions, as indicated by high performance costs in trials where perceptual information conflicted with the endogenous intention (Exp. 1). Additionally, Changes of Intention were more frequent when motor costs of intention pursuit were high (Exp. 2). These findings were simulated using an attractor network model that continuously integrates voluntary intentions, sensory evidence, and motor costs. This argues in favour of a unifying framework for dynamic decision-making processes, in which voluntary actions emerge from a dynamic combination of internal action tendencies and external environmental factors.

## 1 Introduction

People frequently change their minds about what to do. We may plan to go to the gym but end up watching TV, or we may cancel dinner plans after a tiring day at work. Initial action decisions are followed by continuous evaluation processes during which additional information is acquired and integrated with the initial intention. This updating can account for ‘Changes of Mind’ (CoM), i.e., reversals of an initial decision. Most studies investigating CoM focussed on action choices driven by external information (Goodale, Pélisson, & Prablanc, 1986; Resulaj, Kiani, Wolpert, & Shadlen, 2009; Moher & Song, 2014). For example, in the random-dot motion (RDM) task, participants judge the direction of moving dots by reaching for a target that corresponds to the observed dot motion. When the perceptual evidence is noisy, movement trajectories occasionally indicate a CoM, e.g., the response is initiated towards the left, but is then redirected and ends in the right target (Resulaj et al., 2009; Moher & Song, 2014; van den Berg et al., 2016). This suggests that decision making continues after action initiation, allowing for updates of action decisions during ongoing movement execution.

Few studies have considered CoM in the context of voluntary actions. Voluntary actions may be defined as actions in which people endogenously (internally) generate an intention regarding which of several actions to make (Brass & Haggard, 2008).

In many cases, an intention needs to be combined with external sensory input to specify exactly how to move (Schüür & Haggard, 2011). Most previous literature has acknowledged that the way to achieve a goal, or to complete an intentional action, may involve updating based on current circumstances. We propose that the internally-generated intention and external information might both change during the course of action. This possibility suggests two dissociable types of CoM that might occur in voluntary action. In the first type of CoM, external evidence for action may change, as in the RDM task. In the second type of CoM, the intention or goal of action may itself change – a process sometimes called ‘goal-shifting’ (Rubinstein, Meyer, & Evans, 2001). For example, imagine you want to have sushi for dinner (endogenous intention), but you get lost on your way to the Japanese restaurant. After having asked someone for directions (new sensory evidence), you realise you need to generate a new motor command (‘*Change of Movement’*). If the anticipated costs of updating the motor command are high, the initial intention might be changed. For example, you might in the end decide to go to a pizzeria that is just around the corner (‘*Change of Movement and Intention’*) instead of pursuing your initial intention to go to the distant sushi restaurant.

This distinction can be clarified in terms of Pacherie’s hierarchical theory of intention (Pacherie, 2008), Changes of Movement involve changes of ‘motor intentions’, which specify the ‘*how*’ of an action, or its means (i.e., motor policy), rather than its overarching goal. In line with Hebb’s (1949) notion of motor equivalence, there are many ways to achieve a given goal. Hence, movements can be changed while pursuing one and the same initial goal. In contrast, Changes of Intention concern higher-order, distal intentions, which contain conceptual representations of the *goal* or outcome of an action (‘*what*’ decision) (Pacherie, 2008). While updates of motor intentions have been studied extensively (Goodale et al., 1986; Fleming, Mars, Gladwin, & Haggard, 2009; Obhi, Matkovich, & Gilbert, 2009), less is known about how and when people decide to pursue or abandon distal action goals – despite wide-ranging personal and social importance of Changes of Intention (Goschke, 2014). We hypothesised that Changes of Intention depend on the strength of the initial intention and the cost associated with intention pursuit. For instance, a person should be more likely to pursue the intention of having sushi, the stronger that intention is (Ajzen, 1991; Sheeran, Webb, & Gollwitzer, 2005; Fleming et al., 2009). Intentional strength in turn might depend on confidence regarding an internal decision (Folke, Jacobsen, Fleming, & De Martino, 2016) or choice values (Rushworth, 2008), e.g., a strong preference for sushi over pizza. Yet, few goals are worth pursuing at *any* cost. Therefore, time- and effort-related movement costs might induce Changes of Intention, or at least make them more likely to occur (Shadmehr, Orban de Xivry, Xu-Wilson, & Shih, 2010; Cos, Bélanger, & Cisek, 2011).

Both intentional strength and motor costs need to be continuously integrated with momentary sensory evidence to guide action selection. In order to model the neurocognitive mechanisms underlying such dynamic integration of action-relevant sources of information, we propose a unifying computational framework of different types of CoM in voluntary action. Previous computational accounts of CoM have either used bounded accumulator models (Resulaj et al., 2009; Burk, Ingram, Franklin, Shadlen, & Wolpert, 2014; van den Berg et al., 2016) or attractor network models (Albantakis & Deco, 2011; Albantakis, Branzi, Costa, & Deco, 2012; Yan, Zhang, & Wang, 2016; Atiya, Rañó, Prasad, & Wong-Lin, 2019). In both types of models, a decision variable evolves over time given continuous consideration of evidence until a decision threshold is crossed. While accumulator models typically only consider a single source of (sensory) evidence, neural network models represent dynamical systems of interconnected neurons, and thus readily allow for integration of multiple sources of information (Lo & Wang, 2006; Cisek, 2012; Christopoulos, Bonaiuto, & Andersen, 2015). Hence, we here propose an attractor network model that continuously integrates voluntary intentions with sensory evidence and motor costs, accounting for dynamic changes in both lower-level sensorimotor aspects of action selection (Changes of Movement) as well as changes in higher-order voluntary goal intentions (Changes of Intention).

Finally, in addition to changing *objective* action characteristics (e.g., changing movement trajectories), CoM may also shape our ‘Sense of Agency’ (SoA) – the *subjective* experience of exerting control over actions and their outcomes (Pacherie, 2008; Haggard & Tsakiris, 2009). Specifically, changing an ongoing movement could reduce SoA by making actions feel ‘dysfluent’ (Wenke, Fleming, & Haggard, 2010; Sidarus & Haggard, 2016). Whether changing an endogenous intention would affect SoA is less clear. Previous research suggests that strong distal action goals boost SoA (Metcalfe, Eich, & Miele, 2013; Vinding, Pedersen, & Overgaard, 2013). Consequently, deviations from initial intentions might decrease SoA (Villa, Tidoni, Porciello, & Aglioti, 2018). In contrast, reconstructive theories view conscious intentions as retrospective confabulations (Wegner, 2002). People appear to experience actions as intentional, even when they were not part of an initial plan, or are not even their own (Wegner, Sparrow, & Winerman, 2004; Aarts, Custers, & Wegner, 2005). On this view, reversals of endogenous intentions should not affect SoA.

In the current study, participants performed a novel version of the RDM task in which they had to integrate the perceptual decision about dot-motion direction (left/right) with an endogenous choice about which colour to paint the dots. Based on previous studies, we expected to observe perceptual CoM regarding the dot-motion direction (e.g., Resulaj et al., 2009). Importantly, the current paradigm allowed us to differentiate between trials in which perceptual updating about dot-motion direction might produce only updates of the movement (Change of Movement), or might additionally produce an update of the initial colour intention (Change of Movement + Intention).

Importantly, the current paradigm allowed us to differentiate between trials in which perceptual updates only resulted in updates of the movement (Change of Movement), or additionally, an update of the initial colour intention (Change of Movement + Intention). In two experiments, we tested the hypotheses that Changes of Intention are more frequent when initial intentions are weak (Exp. 1) and when the motor cost of intention pursuit is high (Exp. 2). Additionally, in both experiments, subjective reports of SoA were obtained to investigate the effect of CoM on the phenomenology of action. Finally, an Attractor Network Model is introduced that captures dynamic execution, and occasionally reversals, of action decisions.

## 2 Results

### 2.1 Experiment 1: Colour RDM task

Participants performed an adapted version of the RDM task (**Fig. 1**). At the beginning of each trial, they freely chose between two colours (e.g., blue vs. green). Participants were instructed to say the chosen colour in their head, and on 10% of trials, were prompted to say their choice out loud. An RDM stimulus and 4 targets, 2 of each colour, then appeared in pseudo-random locations. Using a touch pad, participants had to move the cursor to the target that matched both the perceived dot-motion direction and their endogenous colour choice (e.g., left-blue target). Once participants reached the target, 25/50/75/100% of dots were presented in the colour of the chosen target and participants were then asked “How much control did you experience over the colour of the dots?” (*SoA judgment*) or “what percentage of dots was painted in the colour you chose?” (% *outcome estimate*).

**Figure 1.**
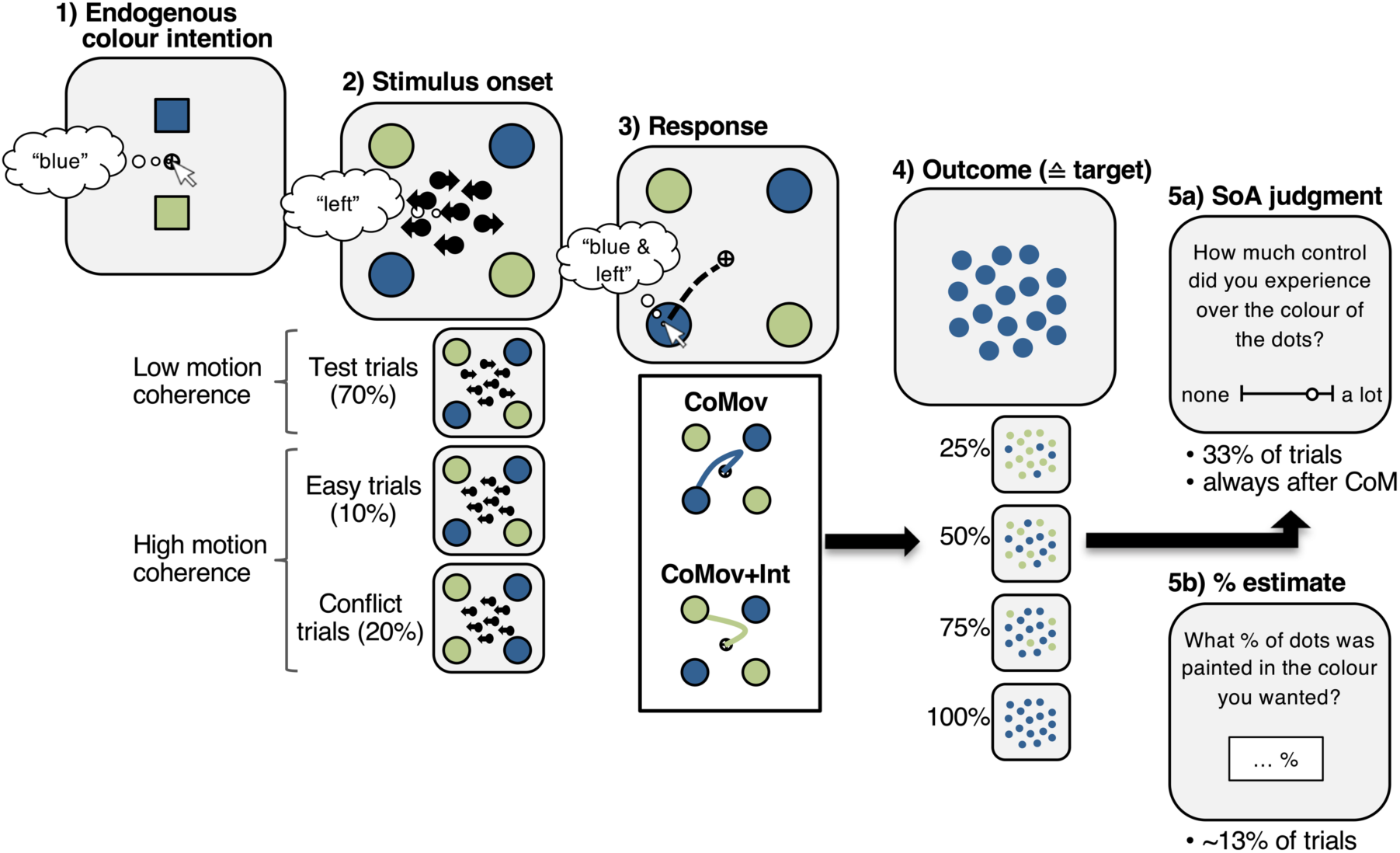
Colour RDM task. Participants generated an endogenous colour intention (1) that had to be integrated with the sensory input of the RDM stimulus (2). Responses were indicated by moving the cursor to the target that matched both the colour intention and dot-motion direction (3). Continuous movement trajectories were measured during response execution allowing for online classification of ‘Changes of Movement’ (CoMov) and ‘Change of Movement + Intention’ (CoMov+Int). Once participants reached the target, 25/50/75/100% of the dots were painted in the colour of the hit target (4). On some trials, participants were asked to provide SoA judgements (5a) or to estimate the percentage of dots that matched their initial colour intention (5b).

In the main condition of interest – *test trials* (70%) – targets of the same colour were presented in diagonally-opposite locations (e.g., blue in top-right and bottom-left corner). Furthermore, motion coherence was low in test trials, with the precise value being determined individually prior to the experiment to ensure ∼60% perceptual choice accuracy (see *Methods*). By contrast, e*asy trials* (10%) served as a baseline condition with high motion coherence (80% coherence). As expected, perceptual choice accuracy was significantly worse in test trials (*M* = 56.6%, *SD* = 9.1%) than in easy trials (*M* = 93.4%, *SD* = 7.0%, *t*(16) = 20.13, *p* < .001, *d* = 4.88) and RTs were significantly slower in test (*M* = 570.5 ms, *SD* = 58.3 ms) than in easy trials (*M* = 534.2 ms, *SD* = 41.5 ms, *t*(16) = 3.99, *p* = .001, *d* = 0.97).

### 2.2 Changes of Mind

Next, we checked whether difficulty of perceptual decisions in test trials resulted in CoM. In analogy to the original RDM task, CoM was defined as a decision reversal regarding the dot-motion direction (e.g., initial response towards a target on the right followed by a switch to a left target). As expected, such perceptual changes were significantly more frequent in test (*M* = 7.64%, *SD* = 6.74%) compared to easy trials (*M* = 1.28%, *SD* = 2.33%; *b* = 1.84, 95% CI [1.19, 2.63], OR = 6.27, χ^2^(1) = 45.69, *p* < .001). More importantly, the current paradigm allowed us to differentiate between trials in which perceptual CoM only resulted in 1) a Change of Movement while the initial colour intention was pursued (*CoMov*; e.g., switch from right-blue to left-blue target), or 2) a Change of Movement that *additionally* involved a Change of Intention (*CoMov+Int*; e.g., switch from right-blue to left-green target). Given the diagonal target arrangement, intention pursuit (CoMov) was associated with longer movement paths than switching to the neighbouring target of different colour (CoMov+Int). Hence, when participants changed their mind about the dot-motion direction, they could save motor costs by switching to the target that did not match their initial colour choice. **Fig. S1** in Supplementary Material shows single-trial movement trajectories for CoMov and CoMov+Int of an individual participant. The average frequency of each type of CoM in test vs. easy trials is shown in **Fig. 2A**.

**Figure 2.**
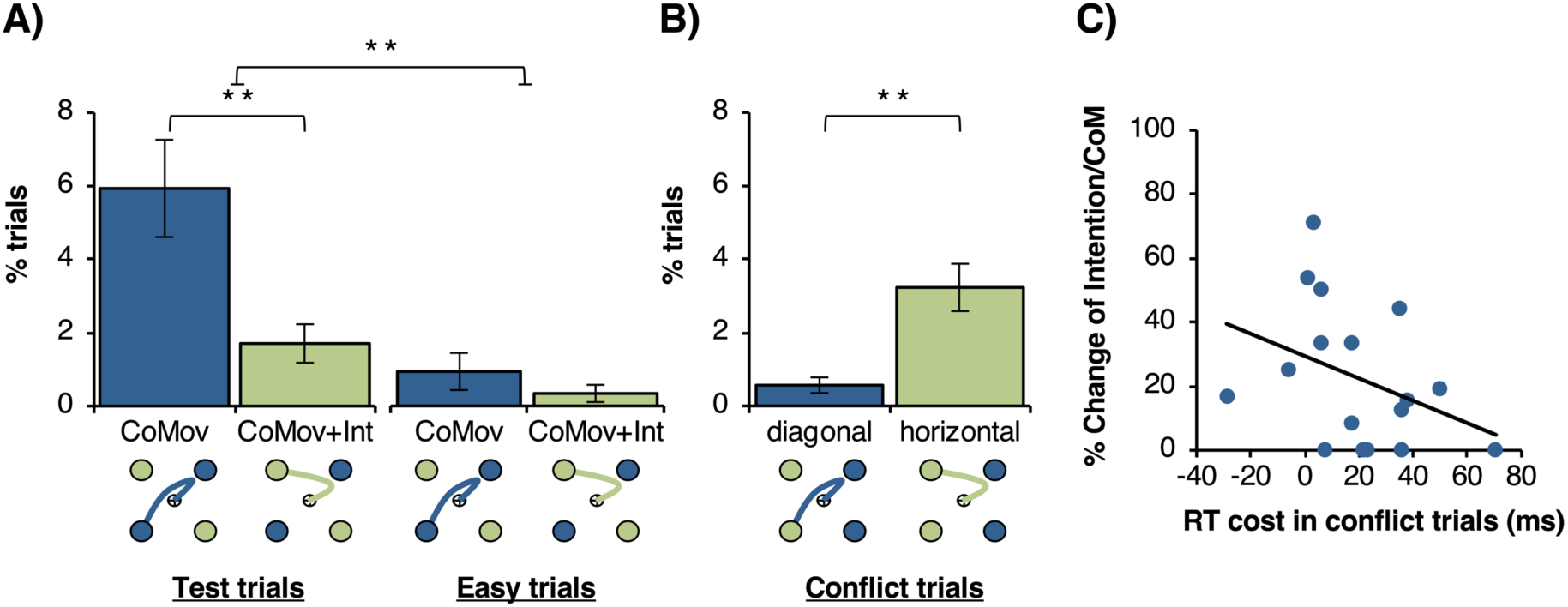
Changes of Mind in the colour RDM task (Exp. 1). A) Percentage of trials classified as ‘Changes of Movement’ (CoMov) and ‘Change of Movement + Intention’ (CoMov+Int) in test and easy trials. B) Percentage of conflict trials with diagonal and horizontal movement corrections of partial errors that were induced by mismatches between colour intentions and dot-motion direction (mean +/- 1 SEM; ** *p* < .01, *N* = 17). C) Correlation across participants between RT costs in conflict trials and frequency of Changes of Intention (relative to overall percentage of CoM).

In order to investigate the relative frequency of CoMov vs. CoMov+Int, only trials with CoM were included in an MELR, with CoMov (0) vs. CoMov+Int (1) as outcome variable and trial condition (easy/test) as a fixed effect. The effect of trial condition was not significant (*b* = −0.13, 95% CI [-1.93, 1.67], OR = 0.88, χ^2^(1) = 0.02, *p* = .890), suggesting that perceptual uncertainty only affected whether or not a perceptual CoM occurred, but did not affect whether participants changed their mind about their colour intention. Interestingly, in test trials, CoMov was more frequent (*M* = 5.9%, *SD* = 5.5%) than CoMov+Int (*M* = 1.7%, *SD* = 2.2%), as indicated by an intercept that was significantly lower than 0 (*b*_0_ = −1.56, 95% CI [-2.47, −0.89], OR = 0.2, *z* = −4.42, *p* < .001). Hence, when changing a movement based on new sensory evidence, participants pursued their colour intention more often than switching to the target of different colour, despite the extra motor costs of diagonal compared to horizontal movement adjustments. A similar trend was observed in easy trials, although overall CoM frequency was low in this condition and the intercept was not significantly different from 0 (*b*_0_ = −1.43, 95% CI [-3.30, 0.43], OR = 0.24, *z* = −1.50, *p* = .132).

#### 2.2.1 Did participants generate initial colour intentions?

As mentioned above, in 90% of trials, participants were *not* asked to verbalise their colour choice at trial start, and instead, colour choices were inferred from movement trajectories (e.g., movements initiated towards green target reflect colour choice of green). This minimised demand characteristics that might discourage participants from changing their initial colour choice when having to say it out loud. Yet, it raises the question whether participants indeed chose a colour at trial start (frame 1, **Fig. 1**), or instead, delayed their decision to stimulus onset (frame 2, **Fig. 1**). The fact that, overall, participants were reluctant to giving up their colour intentions suggests that they assigned a relatively high importance to colour choices in the task. Additionally, we included *conflict trials* (20%) to further investigate whether participants indeed generated colour intentions at trial start, even on trials where they did not have to verbalise their choice. In conflict trials, motion coherence was as high as in easy trials (80% coherence), but both targets of the same colour were on the same side of the screen. Consequently, in roughly 50% of conflict trials, there was a mismatch between intentional colour choice and dot-motion direction (e.g., a participant had chosen blue, both blue targets appeared on the right side, but the dots moved to the left). In this case, participants were instructed to respond according to the dot motion, and hence, move to a target that did not match their own colour choice. If participants did indeed generate initial colour intentions, colour-motion mismatches would induce response conflict. Consequently, RTs and error rates would, on average, be higher in conflict than easy trials even though the perceptual decision was equally easy in both conditions. These performance costs would be driven by trials in which conflict occurred. However, the inference is based on mean performance, and does not require explicitly identifying which specific trials involved conflict and which did not. Note that no response costs would be observed if participants did not make colour choices at trial start since, in that case, participants would simply respond based on dot motion direction without any conflict induced by colour choices.

We found that in conflict trials, RTs were indeed significantly slower (*M* = 549.7 ms, *SD* = 45.8 ms) and perceptual choice accuracy was descriptively lower (*M* = 90.5%, *SD* = 7.2%) than in easy trials (RTs: *M* = 534.2 ms, *SD* = 41.5 ms, *t*(16) = 2.51, *p* = .023, *d* = 0.61; accuracy: *M* = 94.1%, *SD* = 6.8%, *t*(16) = 2.11, *p* = .051, *d* = 0.51). These response costs were present even in trials where targets were presented 700–1000 ms before random-dot stimulus onset (*early targets* in 50% of trials; RT cost: *M*_*Δ*_ = 29.7 ms, *SD*_*Δ*_ = 32.7 ms, *t*(16) = 3.74, *p* = .002, *d* = 0.91; accuracy cost: *M*_*Δ*_ = 3.63%, *SD*_*Δ*_ = 7.27, *t*(16) = 2.06, *p* = .056, *d* = 0.50). This suggests that response costs in conflict trials were not simply driven by participants being surprised about the uncommon target configuration. Instead, participants seemed to generate initial colour intentions, which on some conflict trials did not match the RDM direction, hence inducing response costs.

In addition to errors and slowing of RTs, movement trajectories in conflict trials indicated that participants occasionally initiated a response towards one target, but then adjusted the movement to end in another target, similar to CoM in test/easy trials (**Fig. 2B**) In conflict trials, movement adjustments presumably reflect partial errors in colour-motion mismatch trials. That is, participants initiated responses towards their chosen colour, but then corrected themselves to respond according to the dot motion as instructed. In line with this, we found that corrective movements in conflict trials occurred significantly more often than perceptual CoM in easy trials, despite dot-motion coherence being matched in both conditions (*b* = 1.09, 95% CI [0.39, 1.92], OR = 2.97, χ^2^(1) = 9.93, *p* = .002). This confirms that corrections in conflict trials were not induced by perceptual noise, but instead, can be attributed to conflict induced by mismatches between colour intention and perceptual input.

Finally, in conflict trials, participants could correct ongoing movements in two ways (**Fig. 2B**) by either switching to the diagonally opposite target (similar to CoMov in test trials) or the horizontally neighbouring target (as in CoMov+Int). An MELR showed that participants overall preferred horizontal over diagonal movement corrections in conflict trials (*b*_0_ = 1.74, 95% CI [0.97, 3.24], OR = 5.71, *z* = 3.46, *p* < .001). This suggests that participants were sensitive to the higher motor costs of diagonal movement corrections and preferred less costly horizontal corrections. The fact that, in test trials, participants preferred diagonal (CoMov) over horizontal (CoMov+Int) movements showed that most participants were willing to overcome these motor costs to pursue their colour intentions when possible. However, the frequency of CoMov relative to all CoM in test trials varied across participants (*M* = 77.4%, *SD* = 22.1%). Thus, participants may have differed in how much weight they assigned to the colour choice relative to the perceptual task, and hence, how strong their colour intentions were.

#### 2.2.2 Effect of intentional strength on Changes of Intention

While participants were instructed to generate colour intentions at trial start, they were not explicitly told that they had to *maintain* their initial colour choice throughout the trial. In particular, participants did not receive any instructions as to whether they should stick with their initial colour intention when they changed their mind about the dot-motion direction. Instead, in trials with perceptual CoM, the decision between a) switching to the other target of the same colour or b) switching to the nearby target of different colour was endogenous. This enabled us to capture spontaneous Changes of Intention. Furthermore, the importance of pursuing colour choices was ambiguous on purpose to allow us to capture inter-individual differences in intentional strength – that is, the importance, or weight, a given participant assigned to the colour choice relative to the perceptual choice. For example, a participant who considered colour choices to have little task relevance, would generate weaker intentions, and should be less likely to stick with an initial colour choice when facing the higher cost of colour pursuit in CoM trials.

We tested whether participants with stronger colour intentions showed fewer Changes of Intention. Individuals’ average response costs in conflict compared to easy trials served as an indicator of the strength of colour intention, with higher response costs indicating stronger intentions. Since only 9/17 participants made errors in conflict trials, we focused on RT costs as an indicator of the strength of colour intention. The difference in RTs in conflict–easy trials was correlated with the relative frequency of Changes of Intention out of all 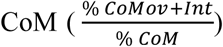 in test trials. As predicted, we found that, across participants, higher RT costs in conflict trials, indicating stronger colour intentions, were indeed associated with fewer Changes of Intention in test trials (Spearman’s ρ(15) = - .50, *p* = .043, 95% CI [-.07, -.76]; **Fig. 2C**). This suggests that participants with stronger colour intentions (and thus higher conflict costs) were less like to change their intentions in test trials.

#### 2.2.3 Potential effect of target confusion

One potential alternative interpretation of trials classified as CoMov+Int needs to be addressed. It is possible that participants switched to a target of different colour because their initial movement was erroneously directed towards a target that did not match their colour choice due to difficulties in target detection. In that case, curved trajectories would not represent a genuine Change of Intention, but rather a correction of an initial colour error. However, a significant number of CoMov+Int was observed even in test trials with early target onset in which participants had 700–1000 ms to identify target-colour locations (*M* = 1.37%, *SD* = 1.85%, *t*(16) = 3.06, *p* = .008, *d* = 0.74). Moreover, participants were rewarded based on perceptual choice only, and hence, switching between horizontal targets merely based on colour would result in a potential monetary loss. Instead, if target confusion occurred, participants should switch to the target of different colour on the same side of the screen (rather than to the horizontally neighbouring target). Importantly, these vertical movement corrections were not classified as CoMov+Int^1^. However, vertical movement corrections were indeed observed on 3.24% of test trials (*SD* = 2.56%) and occurred significantly more often in late-than early-target onset test trials (*b* = 1.18, 95% CI [0.79, 1.59], OR = 3.25, χ2(1) = 38.76, *p* < .001). This suggests that, when participants confused target colours due to difficulties in target detection (e.g., due to late target onset), they switched to the target of different colour that was on the same side of the screen. By contrast, switches to the horizontally neighbouring target (CoMov+Int) presumably represent genuine Changes of Intention that were caused by an initial CoM about the dot-motion direction, rather than target confusion.

### 2.3 Summary & Discussion Exp. 1

In a novel paradigm, two types of CoM in voluntary action were dissociated based on movement trajectories: 1) ‘Changes of Movement’ in which participants changed decisions about exogenous stimuli, requiring them to update motor commands while still pursuing their initial endogenous intention and 2) ‘Changes of Movement + Intention’ where movement updates did not only reflect decision reversals about exogenous stimuli, but additionally, a change of the initial endogenous intention. Although the frequency of CoM was generally, the observed 7.6% CoM in test trials is clearly comparable with previous studies reporting 2–15% CoM in trials with similar motion coherences (Resulaj et al., 2009; Moher & Song, 2014; van den Berg et al., 2016). Further, several areas of cognitive theory, e.g., memory research, rely strongly on data from infrequent errors – no doubt because errors are highly informative about the processes generating performance (Loftus, 2005). Finally, the frequency of CoM varied systematically across trial conditions. Specifically, in line with previous studies on perceptual decision reversals, we found that CoM was more frequent when sensory noise was high, and hence, when initial responses were initiated based on weak sensory evidence. Crucially, we found that the need to update an ongoing movement based on new sensory information occasionally induced a change in the higher-order goal intention regarding colour choice, suggesting that movement reprogramming triggered a re-evaluation of the initial goal itself.

Overall, the frequency of Changes of Intention was lower than that of Changes of Movement, suggesting that, within the context of the current task, the endogenous action goal occupied a primary place in the informational hierarchy, relative to the secondary place occupied by the sensory dot-motion stimulus. However, we further showed that the degree of such prioritisation of endogenous goals over sensory evidence varied across participants. Specifically, the frequency of Changes of Intention was inversely related to the strength of participants’ initial intentions. That is, some participants generated stronger colour intentions as indicated by a high performance cost under endogenous-exogenous conflict. These participants were more likely to pursue their initial intention when adjusting an ongoing movement. Inter-individual differences in intentional strength reflected the importance, or weight, participants assigned to colour choices in the task, relative to the dot-motion judgment. These differences in turn were presumably caused by differences in demand characteristics based on individuals’ interpretation of the instructions (Orne, 1962), or the subjective value participants assigned to the colours (Rushworth, 2008), e.g., based on preferences for certain colours. Note that our design did not allow us to capture variability in intentional strength on a trial-by-trial basis, but rather, the strength of the colour choices throughout the task. However, intentions can vary in strength within people, and it is likely that this would affect the likelihood of a person changing an intention in a given situation (Fleming et al., 2009; Salvaris & Haggard, 2014).

In a second experiment, we manipulated the trade-off between intentions and their associated motor costs on a trial-by-trial basis by varying target distances within participants. We hypothesised that the frequency of Changes of Intention increases when the cost of pursuing the initial intention is high. This would provide more direct evidence that intention reversals can be caused by motor costs associated with intention pursuit. Furthermore, it would establish a means to experimentally induce a higher frequency of Changes of Intention.

### 2.4 Experiment 2: Effect of motor costs on Changes of Intention

The task was identical to Exp. 1 with the following exceptions (**Fig. 3**): Target distance varied on a trial-by-trial basis within participants in order to manipulate the relative motor cost of intention pursuit after a perceptual CoM (**Fig. 3A**). In 50% of trials of each condition, the targets of different colour were far (18°; i.e., far horizontal distance), whereas in the other 50% of trials, the targets of different colour were close (6°; i.e., close horizontal distance). To eliminate visual differences in target detection, the distance of targets from the centre was constant across conditions, i.e., for close horizontal targets, vertical distance was far and vice versa. Importantly, in the far-target condition, path lengths for CoMov and CoMov+Int were roughly equal. Conversely, in the close-target condition, path length was substantially shorter for CoMov+Int (**Fig. 3B**). Hence, in the close-target condition, switching to the target of different colour allowed participants to save motor costs, rendering intention pursuit *relatively* more costly than in the far-target condition. This should increase the frequency of Changes of Intention in the close-compared to the far-target condition. In order to enhance the differences in motor costs between target-distance conditions, the cursor speed was 1.8 times slower than in Exp. 1, increasing overall travel distance of movements. Additionally, target onset was early (700–1000 ms before dot motion onset) in 80% of trials in Exp. 2 in order to reduce the likelihood of target confusion (and hence, initial colour errors).

**Figure 3.**
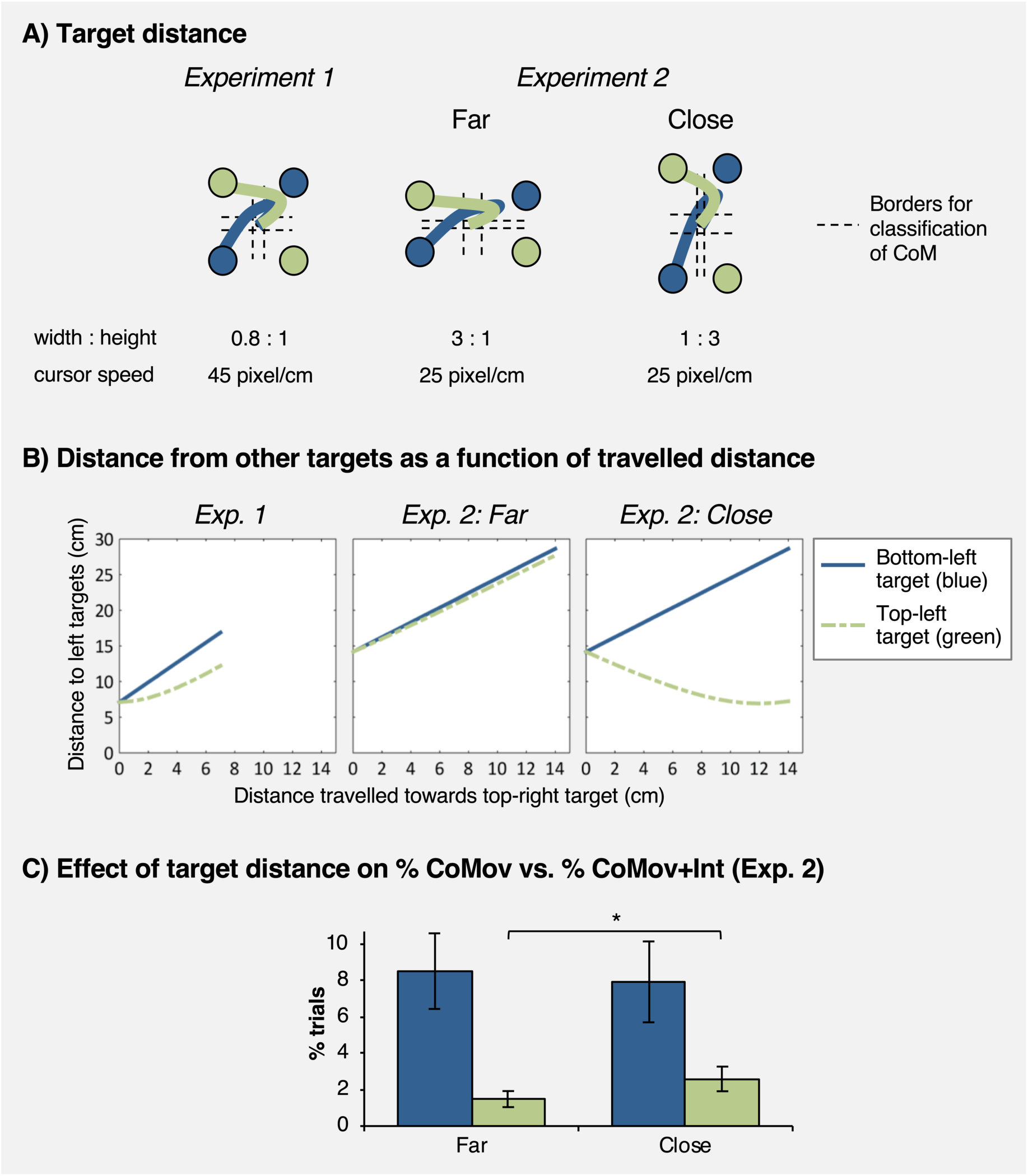
Experiment 2: Manipulation of horizontal target distance. A) Target locations in Experiment 1 and 2. B) Motor costs for each type of CoM as measured by the distance from the diagonal vs. horizontal target as a function of travelled distance (assuming straight movement trajectories towards targets). In the far-target condition, costs associated with each target were roughly equal, whereas in the close-target condition, the target of different colour was closer, hence rendering intention pursuit *relatively* more costly. C) Predicted effect of target distance on frequency of Changes of Intention in Experiment 2.

Overall, behavioural performance in Exp. 2 was comparable to Exp. 1 (see Supplementary Material **S2**). In order to investigate the effect of target distance on the frequency of CoMov vs. CoMov+Int, an MELR model with target distance as a fixed effect (far/close, dummy-coded with far distance as reference level) was conducted for test trials. It revealed a significant effect of target distance (χ^2^(1) = 15.47, *p* < .001), with CoMov+Int occurring more often in the close-(*M* = 2.59%, *SD* = 0.44%) than far-target condition (*M* = 1.48%, *SD* = 0.68%; *b* = 0.76, 95% CI [0.38, 1.16], OR = 2.15, **Fig. 3C**). Interestingly, target distance did not have a significant effect in a model with no-CoM vs. CoM as outcome variable (*b* = 0.06, 95% CI [-0.08, 0.20], OR = 1.06, χ^2^(1) = 0.70, *p* = .404). Hence, target distance did not affect whether or not participants changed their mind about the dot-motion direction, but affected whether or not participants pursued their initial colour choice when a perceptual CoM occurred. That is, cost of goal pursuit was relevant to decisions about goals, but was not relevant to decisions driven by current perceptual input.

Finally, we checked whether the effect of target distance on CoMov+Int depended on target onset time, and hence, the degree to which action cost associated with a potential future change could be anticipated prior to action onset. Including target onset (early/late) in the model revealed no significant main effect of target onset on CoMov+Int vs. CoMov (*b* = −0.05, 95% CI [-0.82, 0.67], OR = 0.95, χ^2^(1) = 0.02, *p* = .902). However, there was a trend for an interaction between target distance and target-onset time (*b* = 0.78, 95% CI [-0.14, 1.74], OR = 2.19, χ^2^(1) = 2.75, *p* = .097). In order to further investigate this interaction, the effect of target distance on CoMov+Int and CoMov was investigated separately for trials with early and late target onset. Relative to no-CoM trials, the likelihood of CoMov+Int increased significantly for close compared to far targets in both the early-target condition (*b* = 0.48, 95% CI [0.12, 0.84], OR = 1.61), χ^2^(1) = 6.95, *p* = .008) and late-target condition (*b* = 0.96, 95% CI [0.33, 1.65], OR = 2.60, χ^2^(1) = 9.10, *p* = .002), with the effect being descriptively stronger in the late-target condition. Additionally, in the late-onset condition, CoMov significantly *decreased* (relative to no-CoM) for close compared to far targets (*b* = −0.46, 95% CI [- 0.83, −0.09], OR = 0.63, χ^2^(1) = 5.94, *p* = .015), which was not the case in early-onset trials (*b* = 0.02, 95% CI [-0.16, 0.19], OR = 1.02, χ^2^(1) = 0.04, *p* = .836). Consequently, the *relative* frequency of Changes of Intention out of all CoM tended to increase more strongly with close targets in the late-target condition than in the early-target condition.

### 2.5 Effect of CoM on SoA (Exp. 1 & 2)

In both experiments, participants were occasionally asked to judge how much control they experienced over the colour of the dots presented at the end of the trial. Participants provided SoA judgments on a visual analogue scale ranging from 0 (“no control”) to 100 (“a lot of control”) after every trial with CoM and in 33% of no-CoM trials. Importantly, after CoM, the percentage of dots painted in the chosen colour was always 50% to avoid that differences in action outcome confound effects of CoM on SoA (for trials without CoM, outcome percentages were assigned randomly for each trial). For analyses of SoA judgments, the data were collapsed across both experiments to increase power (*N* = 33). In test trials without CoM, SoA ratings increased linearly with the percentage of dots matching the colour of the hit target (linear contrast: *F*(1, 32) = 164.91, *p* < .001, η_p_^2^ = .837). When including experiment as a factor, no significant main effect of experiment, nor any interaction were observed (both *F* < 1). Hence, in both experiments, SoA ratings were sensitive to action outcomes showing that participants made appropriate use of the rating scale.

Next, we analysed the effect of CoM on SoA. Variability in trial numbers with CoM was high across participants [*n* CoMov: *M* = 29.8, *SD* = 37.5, range: 1–159; *n* CoMov+Int: *M* = 7.3, *SD* = 9.0, range: 0–43] and 5/33 participants did not show any CoMov+Int. Therefore, linear mixed-effect models were used since they are recommended for analysing unbalanced and missing data (Bagiella, Sloan, & Heitjan, 2000). Furthermore, they allowed us to include continuous predictors that varied on a trial-by-trial level, e.g., movement times. Participants were modelled as random intercepts. For no-CoM, only trials with 50% outcome were included. A model was specified that included CoM as a fixed effect (no-CoM/CoMov/CoMov+Int; dummy coded with no-CoM trials serving as baseline) and SoA ratings as a continuous outcome variable. This model performed significantly better than a model without CoM as a predictor (χ(2) = 13.75, *p* = .001). Post-hoc pairwise comparisons with a Bonferroni-corrected α-level of .05/3 = .017 revealed that the effect of CoM on SoA ratings was driven by a significant decrease of SoA in CoMov (**Fig. 4A**; *M* = 43.8%, *SD* = 9.6%) compared to no-CoM (*M* = 47.1%, *SD* = 8.1%; *b* = −3.02, 95% CI [-4.62, −1.42], *t*(2169.9) = 3.70, *p* < .001), whereas CoMov+Int (*M* = 44.1%, *SD* = 11.0%) did not differ significantly from no-CoM trials (*b* = −1.10, 95% CI [-3.49, 1.29], *t*(2161.2) = 0.91, *p* = .366). The difference between CoMov and CoMov+Int was not significant (*b* = 1.92, 95% CI [-0.48, 1.42], *t*(2162.2) = 1.57, *p* = .118). When adding experiment as a predictor, no main effect of experiment (χ(1) < .01, *p* = .924), nor an interaction with CoM (χ(2) = 0.33, *p* = .847) was found, suggesting that the effect of CoM on SoA was comparable across both experiments.

**Figure 4.**
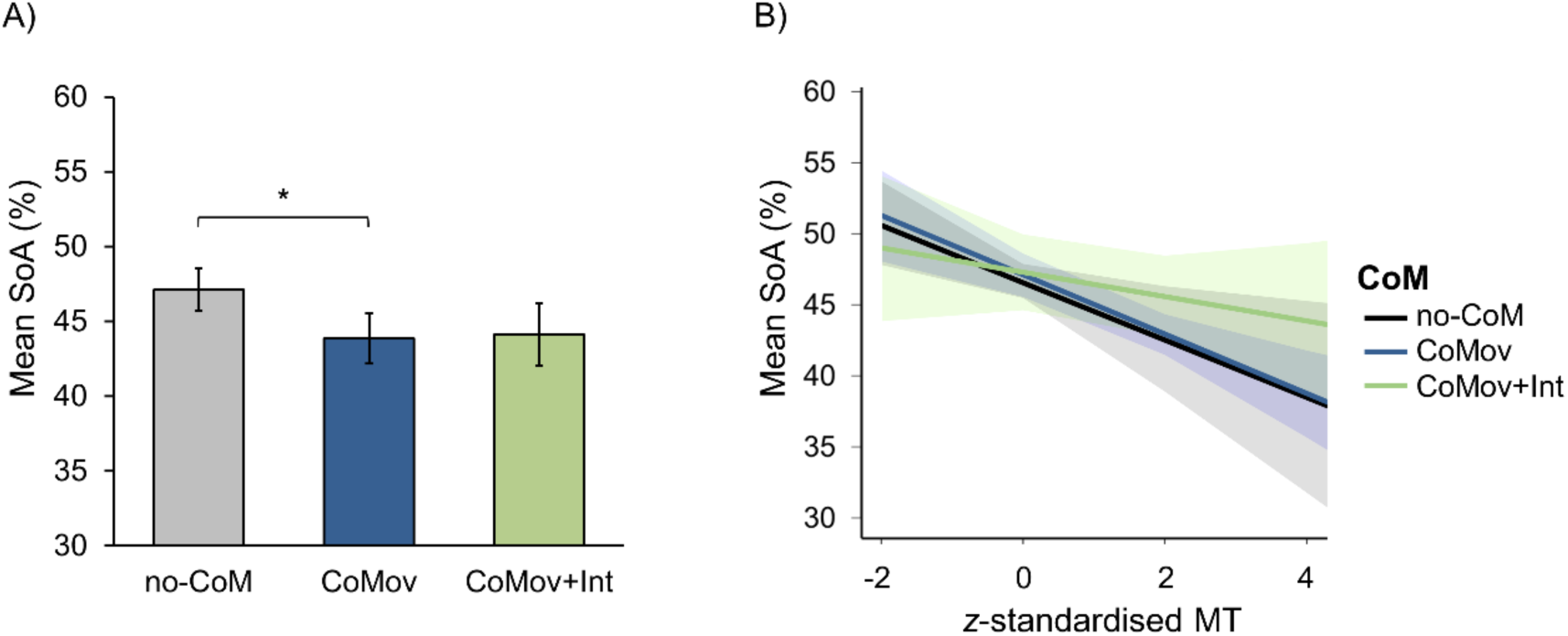
Effect of CoM on SoA (Exp. 1 & 2). A) Mean SoA ratings for each type of CoM. B) Predicted SoA ratings (marginal effects) for a mixed-effects model including movement times (MT) as a predictor (± 1SE, *N* = 33).

As CoM was classified based on movement trajectories, trials differed in terms of pure motor aspects. More specifically, movement times (MTs; i.e., time between response initiation and target hit) were shorter in no-CoM trials (*M* = 480.3 ms, *SD* = 246.8 ms) than in trials with CoMov+Int (*M* = 975.8 ms, *SD* = 365.6 ms, *t*(27) = 10.19, *p* < .001, *d* = 1.93) and CoMov (*M* = 1089.5 ms, *SD* = 354.9 ms, *t*(32) = 18.92, *p* < .001, *d* = 3.29). To investigate if differences in MTs accounted for differences in SoA ratings, individuals’ *z*-standardised MTs were included as a covariate in the model (**Fig. 4B**). This revealed a significant main effect of MTs (χ(1) = 24.32, *p* < .001) driven by lower SoA ratings for longer MTs (*b* = −1.81, 95% CI [-2.58, −1.14]). Furthermore, the effect of CoM on SoA disappeared (χ(2) = 1.51, *p* = .470), and the decrease of SoA ratings in CoMov compared to no-CoM trials was not significant in the model including MTs (*t*(2160.0) = 0.04, *p* = .970). This suggests that the effect of CoMov on SoA was accounted for by differences in MTs. Finally, there was no significant interaction between CoM and MTs (χ(2) = 2.22, *p* = .330). In fact, longer MTs significantly reduced SoA judgments even when only no-CoM trials were considered (*b* = −2.15, 95% CI [-3.58, −0.72], χ(1) = 8.65, *p* = .003), suggesting that MTs affected SoA judgments regardless of whether or not CoM occurred.

### 2.6 Summary & discussion Exp. 2

In Exp. 2, the relative motor cost associated with intention pursuit was manipulated by varying target distances. When the distance to the alternative target colour was short compared to the initially-chosen colour, movement costs for CoMov+Int were low relative to CoMov. This caused an increased frequency of Changes of Intention compared to a condition where targets of both colours were roughly equally distant. Hence, motor costs influenced whether perceptual CoM caused a change in the movement required to realise an intention, or additionally, a change in the intention itself. This effect was present even when targets were presented late (and tended to have a greater effect than when targets were presented early), suggesting that integration of motor costs occurred rapidly and dynamically as actions evolved. That is, even when participants could *not* anticipate action costs before dot-motion onset (late targets), motor costs affected decision making. Thus, action selection does not rely on full anticipation of motor costs (Moher & Song, 2014), but instead, costs may be updated continuously as actions evolve (Todorov & Jordan, 2002; Shadmehr & Krakauer, 2008; Christopoulos et al., 2015).

Finally, across both experiments, reduced SoA was observed after CoMov. However, this effect was statistically accounted for by differences in movement times between trials with and without movement updates. Participants may have used movement times as a proxy of (in)efficient motor performance or difficulty of action selection, which reduces SoA (Wenke et al., 2010; Sidarus & Haggard, 2016). SoA was not modulated when participants changed their initial action goal. These findings are broadly in line with reconstructive accounts of conscious intention, which state that SoA is independent of the actual initial intention and instead relies on retrospective inference (Wegner, 2002; Wegner et al., 2004; Aarts et al., 2005). Indeed, as action goals are updated, predictions about action outcomes may be rapidly adjusted during action (Synofzik, Thier, & Lindner, 2006) without any consequences for subsequent inferences informing SoA. However, the absence of any significant effect of Changes of Intention on SoA in our study should be interpreted with caution since it is a null result based on low trial numbers. In particular, we cannot rule out that strong and sustained intentions may contribute to SoA.

## 3 Attractor network model of CoM in voluntary action

To model the detailed neurocognitive mechanisms underlying CoMov and CoMov+Int, we explored a novel computational approach that could account for the dynamic integration of voluntary intentions with sensory evidence and motor costs. We propose an attractor network model (ANM) that consists of several nodes. Each node represents a population of neurons encoding different modalities of decision-relevant information (**Fig. 5**): 1) neural populations encoding the endogenous colour intention (e.g., blue vs. green; *I*_1_ and *I*_2_), 2) neurons that selectively respond to sensory information about left/right dot-motion (*S*_1_ and *S*_2_), and 3) neurons that calculate the movement cost according to the distance to each of the four target locations (*C*_1_, *C*_2_, *C*_3_, *C*_4_). Information from these neural populations is combined by action nodes (*A*_1_, *A*_2_, *A*_3_, *A*_4_) that integrate all sources of information and specify the motor output, i.e., initiation of a movement towards the chosen target location. For example, action *A*_1_ is selected for execution if the intention is blue (*I*_1_ fires at a high rate), if the dots move left (*S*_1_ fires at a high rate), and if the cost of moving to the left-blue target is relatively low (*C*_1_ fires at a low rate). In other words, the firing rate of each action node reflects the strength of evidence in favour of a given action based on the combined information encoded in a distributed network.

**Figure 5.**
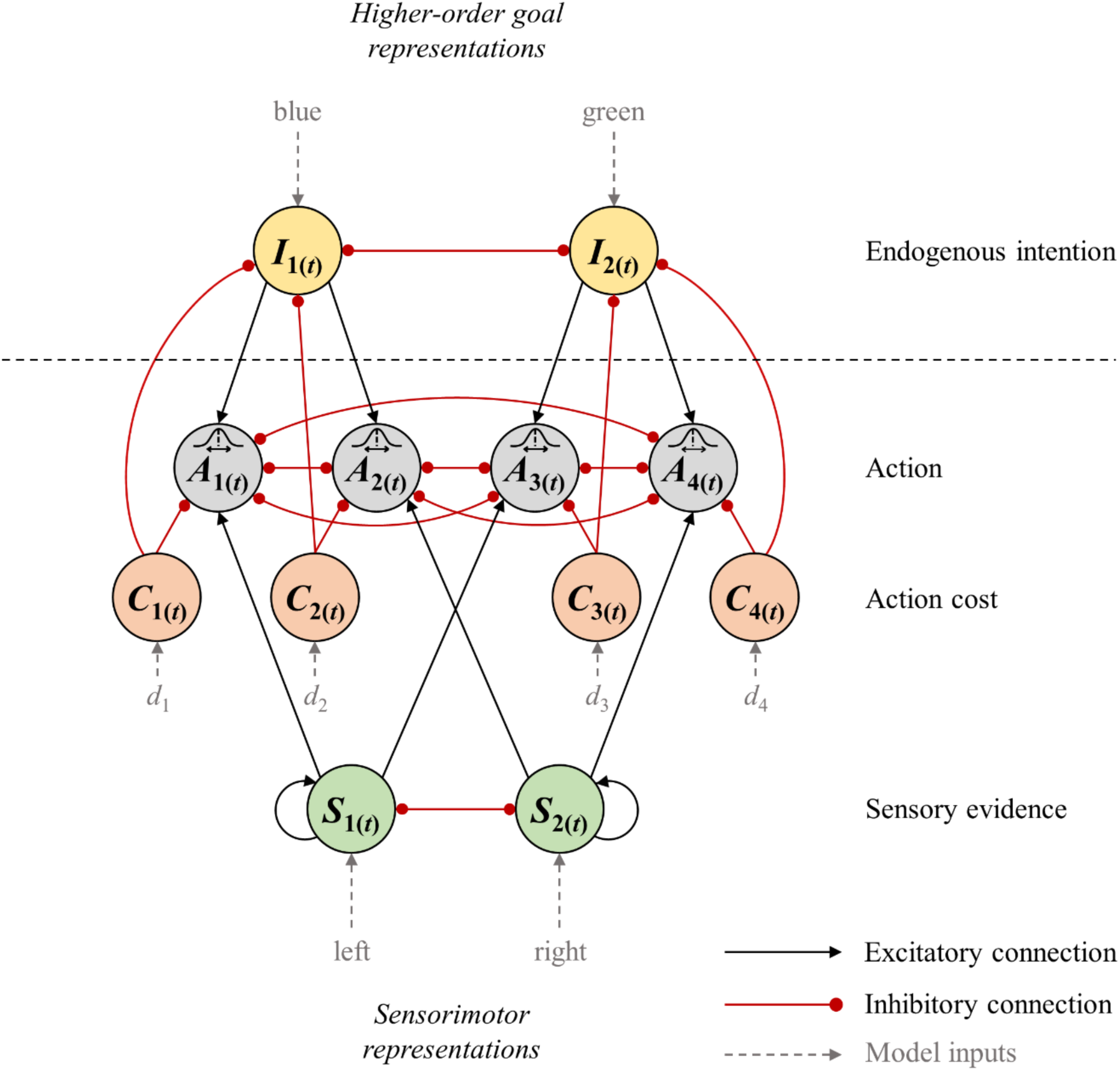
Hierarchical attractor network model of CoM. The network consists of 12 neural nodes that encode different pieces of information. Nodes are connected through excitatory (black) or inhibitory (red) connections. The action nodes *A*_1_ to *A*_4_ compete against each other to determine which one of the four choice targets is selected. This competition takes into account information about 1) endogenous intentions (blue/green represented by nodes *I*_1_ and *I*_2_), 2) sensory information (left/right encoded by sensory nodes *S*_1_ and *S*_2_) and action costs (*C*_1_ to *C*_4_) that depend on the distance *d* to each target location. Intention nodes are represented on a hierarchically higher level than sensorimotor nodes, allowing for top-down regulation of the degree of variability in firing rates of the action nodes. All firing rates are updated continuously and can change dynamically. Hence, CoM can occur when one action node crosses the threshold for movement execution first, but later on, another action wins the competition. Different types of CoM can be dissociated based on which action the network switches to when a decision reversal occurs (e.g., Change of Movement: *A*_1_ → *A*_2_, or Change of Movement + Intention: *A*_1_ → *A*_4_).

The network is further characterised by a hierarchical structure that is based on theories of intention hierarchy (Mele, 1992; Pacherie, 2008; Ondobaka & Bekkering, 2012) and multi-level decision making (Cisek, 2012). Specifically, colour intentions are represented on a hierarchically higher level than sensorimotor information. That is, colour intentions reflect abstract, distal action goals with respect to the action outcome (coloured dots). By contrast, sensorimotor information about perceptual inputs (dot-motion stimulus) and movement-related costs inform *how* that goal can be achieved. Hence, hierarchy in the current model corresponds to the distinction between *what* goal to pursue vs. *how* to pursue it, and hence, the distinction between distal vs. motor intentions (Pacherie, 2008). Hierarchy is implemented as top-down noise regulation in action selection through the intention nodes *I*_1_ and *I*_2_. Specifically, stronger intentions cause a decrease in noise, and thus, decreased variability in firing rates of the action nodes *A*_1_ to *A*_4_. This is in line with previous studies showing that voluntary intentions are associated with noise reduction in motor-related neural activity (Khalighinejad, Schurger, Desantis, Zmigrod, & Haggard, 2018). Additionally, the implementation of hierarchy through noise regulation was inspired by Hierarchical Gaussian Filters, where the degree of noise (or ‘volatility’) of a hierarchically-lower variable can change over time, depending on the current state of a hierarchically-higher variable (Mathys, Daunizeau, Friston, & Stephan, 2011; Mathys et al., 2014). Finally, on each level of the hierarchy, competition between neighbouring network nodes is implemented through lateral inhibition (e.g., *S*_1_ vs. *S*_2_, *I*_1_ vs. *I*_2_, etc.), resulting in a winner-take-all mechanism that determines the final behavioural outcome. Connections across the two hierarchical levels allow for integration of information. Specifically, action representations receive input from higher-order intentions and lower-level sensory evidence as well as information about motor costs. Hence, in the current model, decisions are made through a distributed consensus across different hierarchically-organised neural populations (Cisek, 2012).

Once one of the action nodes reached a fixed firing rate threshold of θ = 40 Hz, a movement towards the corresponding target location was initiated with a motor delay of 180 ms. Crucially, firing rates continued to be updated for 380 ms after initial threshold crossing due to a non-decision time consisting of sensory delays of 200 ms and motor delays of 180 ms (Albantakis & Deco, 2011). This allowed for CoM after action initiation. For example, the model may switch from action *A*_1_ to *A*_2_, reflecting CoMov, i.e., a switch between actions that correspond to different sensory states (*S*_1_ → *S*_2_) but the same colour intention (*I*_1_ → *I*_1_). Alternatively, the network might switch from *A*_1_ to *A*_4_, reflecting CoMov+Int, and hence, a change in both the sensory state (*S*_1_ → *S*_2_) as well as the colour intention (*I*_1_ → *I*_2_). Finally, the model may switch between actions associated with different intentions but the same sensory state (e.g., *A*_1_ → *A*_3_). Note that these types of CoM were not considered in the behavioural task as we assumed that these (vertical) movement switches reflected colour errors where participants erroneously initiated a movement towards a target that did not correspond to their actual initial colour intention (e.g., due to difficulty in target detection). This assumption can be tested in the current model. That is, by defining the ‘true’ colour intention on a given trial, we can analyse to what extent initial colour errors account for CoMov+Int and vertical movement corrections, respectively.

### 3.1 Model implementation & fitting

Details of model implementation and fitting are provided in the *Methods* section. Briefly, firing rates of each neural population were modelled over time using a mean-field approach (Wong & Wang, 2006; Yan et al., 2016). Updates in firing rates depended on 1) how strongly a given node was stimulated (based on external inputs and excitatory/inhibitory inputs from other nodes), 2) the node’s firing rate on the previous time step, and 3) neural noise, which was initially set to 2 Hz in all nodes, but in action nodes, changed over time according to hierarchical noise control through voluntary intentions. The model was fitted to behavioural performance in test trials of Exp. 1. The resulting model was then tested on far/close targets to check whether it could correctly reproduce the effect of target distance on CoMv+Int that we observed in test trials of Exp. 2. Model fitting was performed using a Covariance Matrix Adaptation Evolution Strategy (CMA-ES) algorithm.

### 3.2 CoM in attractor network model

Simulations confirmed that the average outcomes produced by the fitted model closely matched participants’ overall performance in Exp. 1 (see Supplementary Material **S3**). Additionally, we showed that the full model proposed above performed better than alternative models with fewer parameters – e.g., models without cost nodes or without hierarchical noise control (see *Methods*). **Figure S4.1** in Supplementary Material illustrates an example of a single-trial simulation produced by the fitted model. Most importantly, the model produced CoM on some trials (**Fig. 6A**). Average rates of CoMov (*M* = 6.33%) and CoMov+Int (*M* = 1.41%) were highly comparable to CoM observed in Exp. 1 (CoMov: *M* = 5.93%, CoMov+Int: *M* = 1.71%). The model also produced vertical movement corrections on 3.6% of trials (3.24% in Exp. 1). Simulations confirmed that 77.6% of these vertical switches corrected initial colour errors, whereas only 28.4% of CoMov+Int was associated with an initial colour error. Instead, the majority of CoMov+Int produced by the model (56.6%) was associated with a correct initial colour choice, followed by a correction of a *perceptual* error that additionally involved a switch to the alternative colour (CoMov+Int).

**Figure 6.**
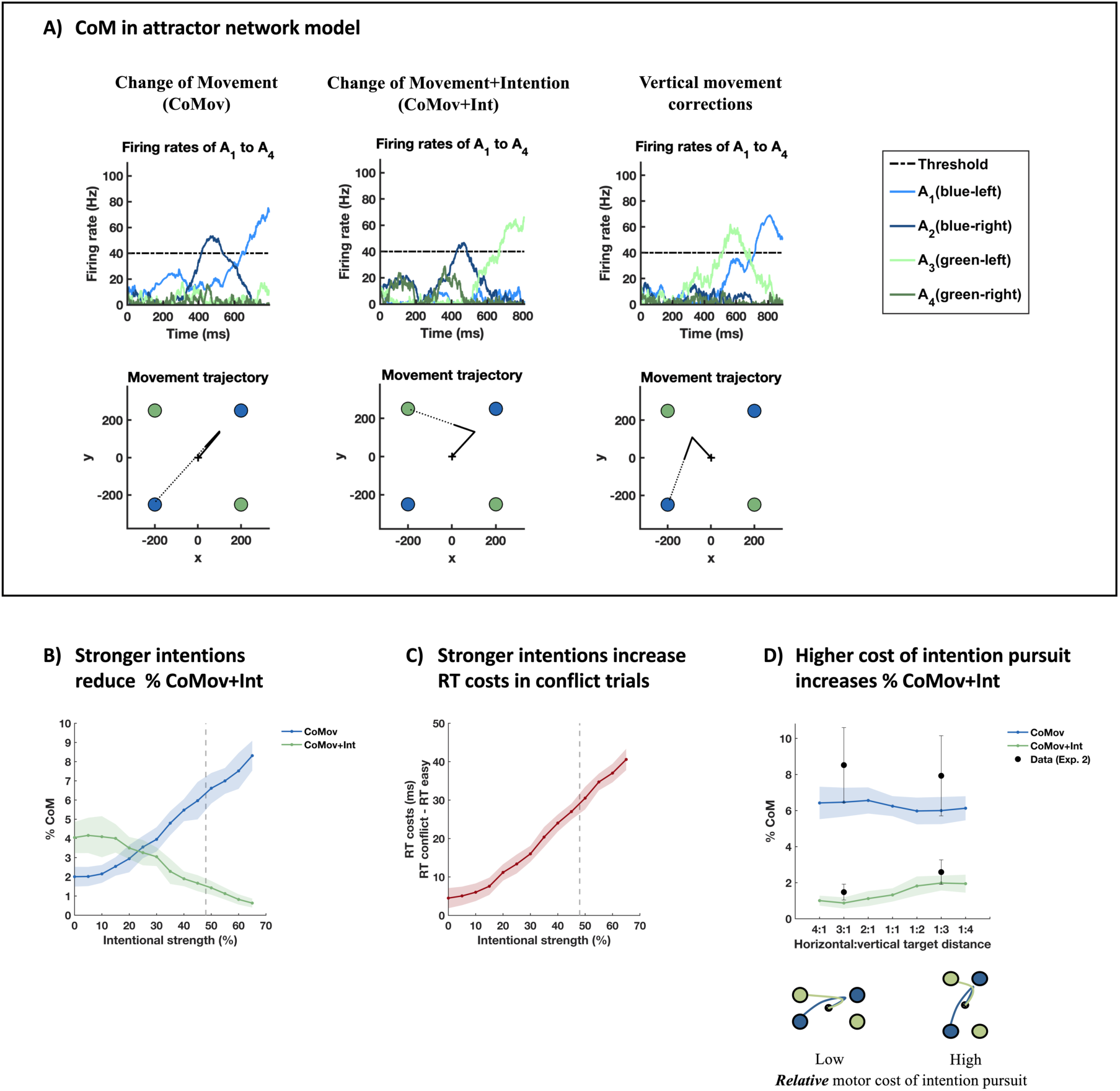
Changes of Mind in Attractor Network Model. A) Single-trial simulations with CoMov, CoMov+Int and vertical movement corrections. Simulated activity of action nodes *A*_1_ to *A*_4_ (top row) and resulting movement trajectories (bottom row). Dotted trajectories indicate completion of movements after the non-decision time of 380 ms (i.e., after the time period during which CoM can occur). B) Effect of intentional strength on CoMov and CoMov+Int. C) Effect of intentional strength on RT costs in conflict trials (relative to easy trials). D) Effect of target locations (and hence, relative motor costs of intention pursuit) on CoMov and CoMov+Int. Grey dashed lines in B) and C) indicate strength of intention in the model that was fitted on data from Exp. 1.

Next, we tested whether the model could reproduce our behavioural findings showing that the frequency of changes in colour intentions depend on intentional strength and motor costs associated with intention pursuit. Indeed, the model was able to capture the effect of intentional strength observed in Exp.1. That is, when increasing the strength of colour intentions while keeping all other model parameters fixed, a decrease in CoMov+Int relative to CoMov was observed (**Fig. 6B**). At the same time, stronger colour intentions also caused the model to produce larger RT costs in conflict trials relative to easy trials (**Fig. 6C**; see Supplementary Material **S5** for details). Thus, in line with our interpretation of the results from Exp. 1, inter-individual differences in the strength of colour intentions may have been the causal factor driving the observed correlation between the frequency of CoMov+Int in test trials and RT costs in conflict trials. Finally, the model fitted to data from Exp. 1 was also able to replicate the effect of target distance observed in Exp. 2. Specifically, when changing horizontal target distance (and hence, the relative cost associated with each action), the model correctly predicted an increase of CoMov+Int in close compared to far horizontal targets. Although this increase was numerically small, it was robust and highly similar to the increase observed in the actual experiment (**Fig. 6D**). The model further predicted a slight decrease in CoMov for far targets, which was descriptively present in Exp. 2. Importantly, we further showed that the effects of strength of intention and motor costs on CoMov+Int in the model were not mediated by potential changes in the rate of initial colour errors. That is, model predictions remained the same when we excluded trials where the model initially chose a target of the wrong colour (see Supplementary Material **S5**).

In summary, the current ANM provides a biologically plausible neuro-computational mechanism through which dynamically changing information from different endogenous and exogenous sources is integrated by a network of neural populations that guide actions in a continuous manner, allowing for rapid Changes of Movement and/or Changes of Intention as actions unfold. Our modelling results confirmed that the majority of CoMov+Int did not reflect corrections of initial colour errors, but instead, reflected true changes in colour intentions that occurred after a correct colour target was initially selected. The frequency of such ‘true’ Changes of Intention in the model in turn depended on the strength of intentions and their associated motor costs, thus replicating the pattern of results observed in the behavioural experiments.

## 4 Discussion

While previous studies of CoM have focused on updating of stimulus-driven actions with arrival of new external evidence (Resulaj et al., 2009; Albantakis & Deco, 2011; Moher & Song, 2014; van den Berg et al., 2016; Atiya et al., 2019), the current study investigated changes of voluntary action decisions. Based on previous research on perceptual CoM, we developed a task where perceptual updates occasionally resulted in an additional change in a higher-order goal intention. In Exp. 1, we showed that the frequency of Changes of Intention was inversely related to the strength of participants’ initial intentions. In Exp. 2, we found that higher motor costs induced more switches to a target that did not match the initial endogenous intention. Finally, we propose that the cognitive mechanisms underlying the flexible nature of voluntary actions can be captured through dynamics in an attractor network model that continuously integrates multiple sources of endogenous and exogenous information.

Past studies of CoM largely neglected such integrative processes, and instead, focused on decisions that are purely driven by a single source of (perceptual) evidence (e.g., Resulaj et al., 2009; Albantakis & Deco, 2011). The current study provides a novel extension of this work by introducing a unified framework for different types of Changes of Mind in voluntary actions, which are guided by several pieces of not just weighted decision variables, but of hierarchically-organised endogenous and exogenous information. Additionally, in contrast to previous computational accounts of CoM, the current model explicitly allowed for an active role of action representations during the evolving decision-making process (Cisek, 2012). That is, action nodes in the current ANM were not simply a mere ‘output’ system of higher-order decision-making areas, but instead, served as a neural hub that integrated different sources of decision-relevant information. In line with previous studies, multiple action representations evolved in parallel, in a dynamic and gradual manner that depended on the current state of evidence in favour of each action, and action selection was determined through a winner-take-all competition between these multiple co-existing affordances (Cisek & Kalaska, 2005; Cisek, 2007; Pastor-Bernier & Cisek, 2011; Selen, Shadlen, & Wolpert, 2012; Thura & Cisek, 2014; Yoo & Hayden, 2018). Moreover, motor outcomes of the model (i.e., simulated movement trajectories) affected subsequent decision updates by causing changes in motor costs (i.e., changes in distance to each target). Thus, our model provides a common framework for decision making and action selection, and accounts for their reciprocal relation, instead of assuming strictly separate and serial processing of decisions and actions (Cisek, 2007; Rushworth, Kolling, Sallet, & Mars, 2012; Yoo & Hayden, 2018). In this context, the model makes further predictions that can be directly tested in future studies. For example, it is plausible that firing rates of action nodes (i.e., the strength of decision evidence in favour of an action) are directly linked to more fine-grained, gradually-varying details of motor policies, such as movement speed or vigour, rather than mere categorical choices between action alternatives.

Finally, the current model implemented a hierarchical organisation between higher-order, abstract intentions and lower-level sensorimotor information. In line with this, previous studies have shown that within the frontal cortex, more anterior regions representing abstract information (e.g., goals) exert top-down control over more posterior regions involved in lower-level sensorimotor control (O’Reilly, 2010; Badre & Nee, 2017). It has been suggested that higher-order areas exert control by ‘gating’ inputs/outputs of lower-level areas (Badre & Nee, 2017). The noise reduction mechanisms implemented in the current model may play an important role in this by enabling gating of action-relevant information, thus allowing intentions to shield action selection from noisy sensory distractions (Kilintari et al., 2018).

We propose that by studying when, why and how voluntary intentions are maintained vs. changed can provide important new insights into the functional role and nature of intentions. Voluntary intentions have previously been conceptualised either as ‘strong determining tendencies’ (Ach, 1935), or instead, as weak and labile plans (Fleming et al., 2009; Salvaris & Haggard, 2014; Kaufman, Churchland, Ryu, & Shenoy, 2015). Rigorous experimental methods to quantify the strength of any given intention have been lacking. Our results suggest that intentions vary gradually in strength, are evaluated continuously, and can be reversed even when an action has already been initiated. Our methods further show that these various features of intention rigidity/flexibility can be quantified and compared within and between individuals. Reversibility of intentions can be highly advantageous in that it allows people to flexibly adjust their behaviour to the current context. On the other hand, an important concept of the voluntary control of behaviour is the need for intention pursuit over long periods of time – e.g., when intending to quit smoking or lose weight. People may give up on these intentions because of new stimuli that can trigger decision reversals. For example, addiction relapse is often caused by exposure to drug-related external stimuli, in particular in individuals with high sensitivity to incentive cues (Robinson, Robinson, & Berridge, 2013). More generally, disturbances in the balance between goal-shielding vs. goal-switching may be linked to a large range of psychiatric and neurological conditions, and hence, understanding the processes underlying this balance is crucial to well-being and mental health (Goschke, 2014).

In conclusion, voluntary actions are shaped by continuous decision-making processes that integrate external information with endogenous intentions. The flexible nature of action selection allows agents to dynamically decide *which* intention to pursue and *how* to pursue it. Our study introduces a quantitative laboratory approach and novel computational model that can capture the neurocognitive processes underlying flexible goal-directed actions, providing important new insights into the nature of voluntary intentions, and the mechanisms underlying goal pursuit and its disturbances, with important social and personal implications.

## 5 Methods

### 5.1 Participants

The study was approved by the UCL Research Ethics Committee. Participants provided written informed consent prior to the study. For Exp. 1, twenty-one right-handed participants were recruited through the ICN subject database. One participant did not reach the performance criterion in the training session (see below) and another participant withdrew after training. Two further participants were excluded, one due to technical issues during data collection and one due to strategic decision delay in the task (see below). The final sample consisted of 17 participants (13 female, age: *M* = 22.6 yr, *SD* = 3.1). For Exp. 2, twenty-one right-handed participants were initially invited for the experiment. Three participants did not reach the performance criterion during training (see below) and two further participants were excluded due to poor performance in the test session (> 15% errors or misses in easy trials), resulting in a final sample of 16 participants (11 female, age: *M* = 23.2 yr, *SD* = 2.9). Participants received £7.50/hour and a performance-based reward.

### 5.2 Apparatus and stimuli

The experiment was programmed in Matlab R2014a and the Psychophysics Toolbox (Brainard, 1997). RDM stimuli were generated using the Variable Coherence RDM code (https://shadlenlab.columbia.edu/resources/VCRDM.html). The stimuli were presented in a central aperture (4.5° diameter) with a stimulus density of 15.6 dots deg^-2^ s^-1^, at a screen refresh rate of 60 Hz. The percentage of dots that were displaced in the same direction determined the motion coherence and motion direction (left/right) was assigned randomly for each trial. Target circles of 1.8° diameter were located at a distance of 9.6° from the centre of the screen (x = 6.0°, y = 7.5°). Target colours were random pairs of blue, green, pink, and orange of comparable luminance. Participants were seated approximately 60 cm from a computer screen and moved a cursor to the targets using a Wacom Intuos Pro pen tablet. Movement trajectories were recorded at a sampling frequency of 125 Hz.

### 5.3 Trial procedure

Participants made endogenous choices between random pairs of 4 target colours (blue/green/pink/orange). Once they had chosen a colour, participants clicked on a central fixation cross and after a random delay of 700–1000 ms, the RDM stimulus and 4 targets, 2 of each colour, appeared. Targets were either presented immediately after colour choice (*early targets*), or they appeared 700–1000 ms after colour choice, i.e., at the same time as the dot-motion stimulus (*late targets*). 500 ms after participants reached a target, 25/50/75/100% of the dots from the last 3 video frames were presented in the colour of the hit target (1 sec). On 1/3 of trials, and after every CoM, participants were then asked to provide SoA judgements on a visual analogue scale ranging from “none” to “a lot”. On 1/5 of the remaining trials (∼13% of trials overall), participants were asked to provide an estimate of the percentage of dots that matched their initial colour intention. Note that outcome judgments were included to motivate participants to pay attention to the action outcomes, and hence, render colour choices more meaningful within the context of the task. However, given that outcome judgments never appeared after CoM (which was always followed by SoA ratings), we did not further analyse them.

### 5.4 Training session

Participants had to pass a training session the day before the actual experiment. They were trained on the original 2-choice RDM task until they reached 70% accuracy in trials with 35% motion coherence. One participant failed to reach the criterion and was not invited for the experimental session. All other participants performed 160 additional trials with randomly varying motion strength (5–65% coherence) in order to obtain stable performance. Finally, an alternating staircase procedure was administered (see Moher & Song, 2014 for details) to determine the motion coherence at which a participant’s accuracy was ∼60% (coherence: *M* = 11.8% *SD* = 4.1%). This level was chosen to maximize the frequency of perceptual CoM (Resulaj et al., 2009). During training, trial-by-trial error feedback (red dots) was provided.

### 5.5 Experimental session

After a short practice block, participants were given 1h to complete as many trials as possible (*M* = 358.2, *SD* = 37.5) in Experiment 1. In Exp. 2, participants completed two identical experimental sessions in which they were given 1.15h each to complete as many trials as (*M* = 815.6, *SD* = 57.2). The duration of Exp. 2 was increased compared to Exp. 1 in order to obtain a sufficient number of CoMov and CoMov+Int for each target distance condition. To motivate participants to be fast and accurate, they won 1 p for every correct perceptual choice. After each block of 30 trials, participants received feedback about their perceptual choice accuracy. There was no trial-by-trial error feedback, but a “too slow!” message was shown if response initiation exceeded a certain deadline or if the target was not reached within 3 sec after response initiation. In order to induce fast response initiation, the response deadline was initially 1000 ms, but decreased by 50 ms after every block if a participant had less than 10% trials with CoM and less than 15% misses. Reaction times (RTs) were defined as the point in time at which the cursor left a central circle of 1.1° diameter, at which point the RDM stimulus disappeared. Previous studies showed that, due to sensorimotor delays, CoM occurs even when the external stimulus is removed at action onset (Resulaj et al., 2009).

In both the training and test session, participants were instructed to fixate the central cross throughout each trial. Electrooculography was used to monitor eye movements and, whenever necessary, participants were reminded to keep fixation.

### 5.6 Classification of CoM

Trials with CoM were classified online based on movement trajectories: If the cursor position exceeded 10% of both the x- and y-distance towards a given target, but then ended in the diagonally opposite target (of the same colour), the trial was classified as CoMov. If it ended in the horizontally neighbouring target (of different colour), it was classified as a CoMov+Int. In Exp. 2. the absolute coordinates that had to be exceeded differed between target distance conditions (**Fig. 3A**), due to the different target locations. Using relative rather than absolute coordinates in Exp. 2 ensured that CoM classification was not biased by differences in movement angles across target distance conditions.

### 5.7 Movement analysis

Movement trajectories were analysed in Matlab R2014b. All trials that had been classified as CoM during the task were inspected individually. Trials with double CoM (Exp. 1: 0.93% of all trials; Exp. 2: 0.83%) or initial movement trajectories that were not clearly directed towards one of the targets (e.g., circular trajectories or vertical movement initiation; Exp. 1: 0.13%; Exp. 2: 0.47%) were excluded from all analyses. Furthermore, velocity profiles of reaching movements were analysed. Note that participants might have initiated a response in any direction in order to comply with the short response deadline, subsequently choosing a target only after having left the home position. In that case, curvature away from the initial trajectory would not be a CoM, as the initial trajectory would not reflect commitment to a specific target. Completely excluding any element of strategic delay for individual trials is difficult. However, frequent stopping shortly after movement initiation (i.e., velocity = 0 at some point *after* movement initiation) even in trials with straight trajectories would clearly indicate strategic decision delay. In Exp. 1, one participant stopped in 28.6% of straight trajectories, with an average stop duration of 351.2 ms and was therefore excluded from all analyses. Such stopping was rare in all other participants (stop frequency: *M* = 7.4%^2^, *SD* = 5.2%; stop duration: *M* = 157.9 ms, *SD* = 40.8 ms). In Exp. 2, movement velocities indicated that none of the participants showed strategic decision delay.

### 5.8 Statistical analyses

Given the small percentage of trials with CoM, mixed-effects logistic regression (MELR) models were used for analyses of CoM frequency (Bagiella et al., 2000). Model fitting was performed using Maximum-likelihood estimation with the lme4 package (Bates, Maechler, Bolker, & Walker, 2014) in R (R Development Core Team, 2015). Binomial models with a logit link were specified. To investigate CoM, two types of binary outcome variables were analysed: Either no-CoM (0) vs. CoM (1) for analyses of overall frequency of perceptual CoM (regardless of type of CoM), or CoMov (0) vs. CoMov+Int (1) for analyses of different types of Change of Mind within CoM trials. Participants were modelled as random intercepts. Including random slopes did not change any of the results and only one of the models performed significantly better when random slopes were added. Hence, all models reported contain random intercepts only. Parameter estimates *b* and 95% profile confidence intervals are reported in log-odds space, and odds ratios (OR) are reported to facilitate interpretation. Statistical inference was performed by comparing models with vs. without a given fixed effect using likelihood-ratio tests. Satterthwaite approximation for degrees of freedom was used (Kuznetsova, Brockhoff, & Christensen, 2015). All other analyses (comparison of means with ANOVAs/*t* tests) were performed in IBM SPSS Statistics for Windows, version 21 (Corp, Released 2012). For RT analyses, only correct trials within +/- 3 SD of the individual’s average RT in each condition were included.

### 5.9 Attractor network model

The model was implemented and fitted in Python 3.7. All model code is available on GitHub, including a Matlab implementation of the model that can be used to run simulations and plot single-trial model outcomes (https://github.com/AnneLoffler/AttractorNetwork-CoM).

#### 5.9.1 Network architecture

The attractor network model consists of 12 neural nodes that are grouped into different modules according to the source of information they represent (**Fig. 5**):

1. Two ***intention nodes*** (*I*_1_, *I*_2_) that encode the voluntary intention (blue/green)
2. Two ***sensory nodes*** (*S*_1_, *S*_2_) that selectively respond to dot-motion direction (left/right)
3. Four ***cost nodes*** (*C*_1_, *C*_2_, *C*_3_, *C*_4_) that calculate the cost associated with each action based on distance to each target location
4. Four ***action nodes*** (*A*_1_, *A*_2_, *A*_3_, *A*_4_) that correspond to the 4 possible action alternatives, and hence, location of the choice targets (left/right top/bottom)

Each node represents a population of neurons whose firing rates change dynamically over time. The firing rates of intention nodes (*I*_1_, *I*_2_) and sensory nodes (*S*_1_, *S*_2_) depend on model inputs whose intensities corresponded to the strength of intention and strength of sensory evidence (i.e., motion coherence), respectively. Firing rates of cost nodes (*C*1 to *C*4) depend on the distance *d* to each target location. Hence, intention, sensory and cost nodes are ‘input nodes’ that receive direct model inputs. Action nodes do not receive any direct external inputs, but instead, integrate information from all other network nodes to determine the behavioural outcome (i.e., movement trajectory towards one of the four targets). Integration of information is achieved through neural connectivity. Colour intentions and sensory inputs have excitatory effects on action nodes, whereas costs have inhibitory effects. Furthermore, neurons that encode the same modality of information, e.g., sensory nodes *S*_1_ and *S*_2_, but respond selectively to a specific input (e.g., left vs. right dot motion), compete against each other through lateral inhibition. This ensures that over time, a single choice option is selected through a winner-take-all mechanism that supresses competing choice alternatives.

Due to considerations of parsimony, network connections were assumed to be symmetric. For example, *I*_1_ and *I*_2_ had equally strong connections onto their corresponding action nodes (and similar for *S*_1_ and *S*_2_ etc.). Additionally, inhibitory competition between action nodes associated with the same colour intention (i.e., *A*_1_ vs. *A*_2_ and *A*_3_ vs. *A*_4_) was assumed to be stronger than competition between action nodes associated with different colour intentions (e.g., *A*_1_ vs. *A*_3_). This was because actions associated with the same colour intention corresponded to diagonally-opposite targets, respectively, and hence, movements in either direction were mutually exclusive (i.e., competition is stronger). Moreover, the effects of costs on intention nodes were assumed to be weaker than the effects of costs on action nodes. Note that this assumption was necessary since action costs would otherwise completely suppress intentions at trial start. Due to considerations of parsimony, we chose to fix the weight from costs to intention nodes to be 0.5 of the weight from costs to action nodes, instead of fitting two separate weights for each cost parameter. Finally, sensory nodes had self-excitatory connections, representing temporal integration of sensory evidence from the dot-motion stimulus (Wong & Wang, 2006; Albantakis & Deco, 2011).

#### 5.9.2 Modelling firing rates

In order to compute the firing rates of each neural node over time, a mean-field approach was used (Wong & Wang, 2006; Yan et al., 2016). That is, instead of modelling individual spiking neurons, the overall firing rate of a given neural population (node) was calculated for each point in time. Firing rates of each node were updated in time steps of 1 ms and depended on 1) how strongly a given node was stimulated (based on external model inputs and excitatory/inhibitory inputs from other nodes), 2) the node’s firing rate on the previous time step, and 3) neural noise. Hence, the following equations were used to determine the firing rate *r* of a given node *i* at time point *t*.

First, the total stimulation that each node received at time *t* was calculated according to equation 1. Stimulation depended on direct external inputs into that node (if any), plus the sum of neural inputs from all other nodes (and itself in case of auto-connections). The neural input that node *i* received from node *j* depended on the firing rate of *j* at the previous time step weighted by its connectivity to *i* as defined by the weight matrix *W*:

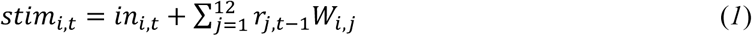

Updates in firing rates were then computed as a function of the node’s previous firing rate, the current stimulation *stim*_*i,t*_ it received, and a base time constant τ of 100 ms (Wong & Wang, 2006; Yan et al., 2016). Hence, using the Euler-Maruyama approximation for differential equations (Miller, 2016; Hahne et al., 2017), the firing rate of node *i* at time *t* was calculated as follows:

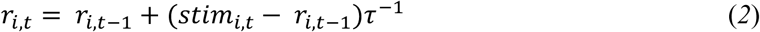

Finally, random Gaussian noise *s* was added to the firing rate of each node:

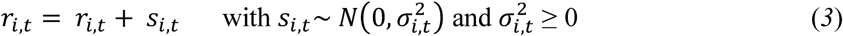

The degree of neural noise varied according to σ^2^, which was initially set to 2 Hz for all nodes. However, according to the assumption of top-down noise regulation through higher-order intentions, σ^2^ of each action node *A*_1_ to *A*_4_ varied as a function of the state of intention nodes *I*_1_ and *I*_2_. Specifically, higher firing rates of *I*_1_ caused a reduction of noise in its associated action nodes *A*_1_ and *A*_2_ (equation 4a) and *I*_2_ caused noise reduction in action nodes *A*_3_ and *A*_4_ (equation 4b) where *h* is a factor that indicates the degree of this hierarchical noise reduction. For example, if *h* = 1 and *I*_1_ fires at 50% of its maximum firing rate, noise in *A*_1_ and *A*_2_ is reduced by 50%:

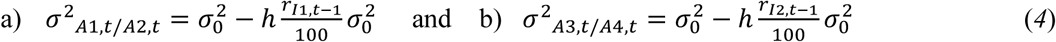

Firing rates were restricted to a range of 0–100 Hz. All neurons started with a background firing rate of 10 Hz. Once one of the action nodes reached a fixed firing rate threshold of θ = 40 Hz (and surpassed all other action nodes by at least 10 Hz to ensure a single winning action), a movement was initiated with a motor delay of 180 ms. Movement direction corresponded to the chosen target location and movement speed was constant at 0.7 pixels/ms, resulting in a movement duration of ∼450 ms for straight movement trajectories, in line with movement times measured in Exp. 1. Movement execution towards a chosen target continued even if the action node dropped below the threshold, unless another action node reached the firing rate threshold, in which case the movement was redirected towards the new target choice. Firing rates continued to be updated for 380 ms after initial threshold crossing due to a non-decision time consisting of sensory delays of 200 ms and motor delays of 180 ms (Albantakis & Deco, 2011). This caused decisions to be updated even after initial action onset (Resulaj et al., 2009; Albantakis & Deco, 2011). After the non-decision time, firing rate updates were stopped and the movement was completed according to the final target choice.

#### 5.9.3 Model inputs

External model inputs were simulated at a rate of *f*_*in*_ = 60 Hz. Inputs into sensory nodes were presented after the sensory delay of 200 ms. The respective strength of inputs into *S*_1_ and *S*_2_ corresponded to the strength of sensory evidence, i.e., dot-motion coherence, and was calculated as:

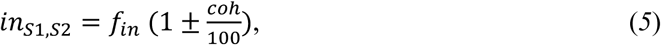

with *coh* corresponding to the % coherence and +/– indicating whether or not motion direction corresponded to the neurons’ preferred motion direction. By analogy, inputs into intention nodes depended on the strength of the endogenous colour intention *col*, i.e., the relative ‘endogenous evidence’ in favour of a given colour:

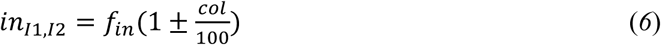

Equations 5 and 6 ensured that model inputs were normalised. Hence, the total input into the network was constant across different levels of sensory/endogenous strength. Similarly, the input into cost nodes was set to an equal value of 60 Hz at trial start. Once a movement was initiated, costs were updated relative to changes in Euclidean distance between the current position and each target location. Consequently, the total external model inputs (sensory, endogenous, costs) were balanced, and thus, only the *relative* strength of evidence from each source affected action selection.

#### 5.9.4 Model fitting

Eight model parameters were fitted, relating to the connectivity weights between neural nodes, the strength of decision evidence, and the degree of top-down noise modulation through intentions:

##### Network connectivity weights

1. Intentional nodes to action nodes (e.g., 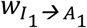)
2. Sensory nodes to action nodes (e.g., 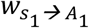)
3. Cost nodes to action nodes (e.g., 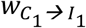) and intention nodes (e.g., 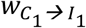)
4. Sensory auto-connections (e.g., 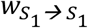)
5. Lateral inhibition (e.g., 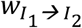)

##### Decision evidence

6) Strength of sensory evidence (dot motion coherence)
7) Intentional strength (colour intention)

##### Hierarchical control

8) Degree of hierarchical noise control

To reduce the number of free parameters, the strength of connectivity weights was assumed to be symmetrical (e.g., 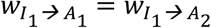). A covariance matrix adaptation evolution strategy (CMA-ES) algorithm with maximum likelihood updates was used to fit model parameters (Hansen & Ostermeier, 1996; Jancke, Igel, & Erlhagen, 2001). CMA-ES is a randomized search algorithm suited to explore our parameter space about which initial knowledge is constrained. We first informally explored different sets of parameters to approximate values that would yield reasonable behavioural outcomes (i.e., RTs within 500-1000 ms, perceptual error rates of ∼40%, CoM of ∼10% with more CoMov than CoMov+Int, and low rates of missed/early responses). We then performed a global optimization test with single- and multi-objective CMA-ES, which validated that one of the convergence regions was on the intuitively preferable parameter values. Finally, this convergence region was used to initialize a set of local optimization runs to fine-tune parameters. A multi-objective CMA-ES was applied (Voß, Hansen, & Igel, 2010; Hansen, 2011) to minimize the error of model predictions from actual behavioural outcomes in Exp.1. Specifically, the model was optimized according to the following behavioural outcomes: RTs, % perceptual choice accuracy, % CoMov, % CoMov+Int, and % colour changes that occurred without a perceptual change (vertical switches). Additionally, the model was fitted to minimise early responses (i.e., responses before stimulus onset), misses (RTs > 1 sec), and maximize colour choice accuracy (e.g., the true intention was ‘blue’ and an action node associated with ‘blue’ was selected). Note that for colour accuracy, only initial choices were considered since final colour ‘errors’ after a CoM may reflect CoMov+Int, and hence, a change in the colour intention rather than an error with respect to the initially chosen target.

Each optimization run consisted of 10000 trials. Powell’s method was used to obtain local minima (Powell, 1964). Table S4.1 summarises the initial parameter values and step sizes that were used for the optimization as well as the final, fine-tuned parameters obtained with the CMA-ES approach. Using the fitted model parameters, 30 simulations of 1000 trials each were then conducted to derive model predictions for a given experimental condition (e.g., simulation of effect of target distance, see section S6.3). For Exp. 1, model predictions strongly overlapped with participants’ actual behaviour in Exp. 1 (**Table S3.2** in Supplementary Material).

#### 5.9.5 Model comparisons

The model obtained through the CMA-ES procedure yielded a good fit to the behaviour observed in test trials of Exp. 1. To check whether this model performed better than simpler versions of the network, we compared the full model to two alternative models with fewer parameters: 1) a model without hierarchical noise control (i.e., when *h* was fixed to 0) and 2) a model without action cost parameter (i.e., when *w*_*C*→*A*_ was fixed to 0). The Akaike Information Criterion (AIC) was used to evaluate model fits, and the relative likelihood of the respective models was then obtained as follows:

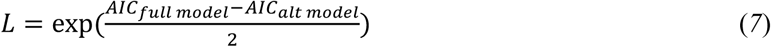

Model comparisons revealed that the model without hierarchical noise control was only 0.139 times as probable (AIC = 35.58) as the full model with 8 parameters (AIC = 31.64). Furthermore, the model without an action costs was only 0.0005 times as likely (AIC = 47.05) as the full model. Thus, the attractor network model we proposed here (**Fig. 5**) was the model that best accounted for the behavioural results observed in Exp. 1.

## Acknowledgments

This work was supported by the European Research Council (grant number 323943). The authors thank Lucie Charles and Eoin Travers for helpful comments on a previous draft of this manuscript.

## Author contributions

A.L. and P.H. conceived and designed the experiments. A.L. performed data collection and analysis. A.L., A.S., and Z.F. implemented the computational model and performed model simulations and fitting. A.L. drafted the manuscript and all other authors provided crucial feedback and revisions. All authors approved the final version of the manuscript.

## Footnotes

1 Note that there was no reason for participants to change their mind about the target colour unless they had changed their mind about the dot-motion direction causing a switch between left/right targets. Consequently, switches between the two targets of different colour on the *same side* of the screen (i.e., vertical switches) were not considered real Changes of Intention within the context of the current task (see Fig. 6a for examples of vertical switches produced by the model).

2 Note that this percentage is highly comparable to the percentage of trials with CoM, and hence, can be attributed to decision uncertainty and vacillation, rather than strategic decision delay.

