## Supplementary Material for "Two Ways to Change Your Mind: Effects of Intentional Strength and Motor Costs on Changes of Intention"

#### S1 Single-trial movement trajectories in test trials

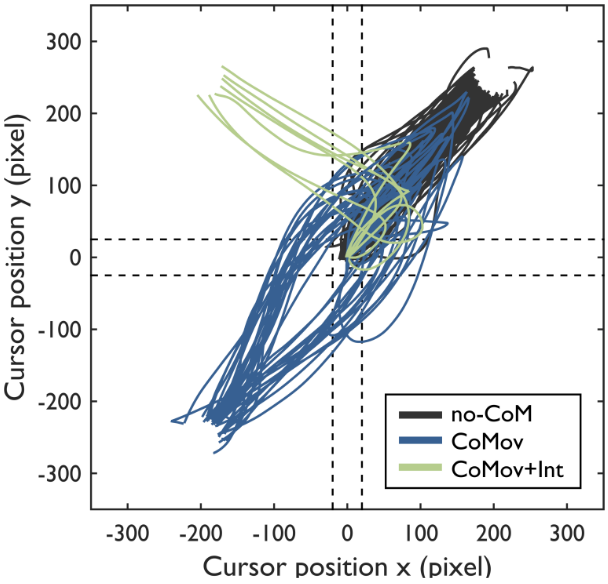


**Figure S1.1**. Single-trial movement trajectories in test trials of one participant. For illustration purposes, trajectories were mirrored such that the initial movement was always directed towards the upper right target, but ended in a different target depending on the class of movement trajectory (no Change of Mind, Change of Movement, or Change of Movement + Intention). Dashed lines indicate the coordinates that were used as criteria for CoM classification.

#### S2 Task performance in Experiment 2

As in Experiment 1, accuracy of the perceptual choice was significantly lower in test trials ($M$ = 58.5%, $SD$ = 5.2%) compared to easy trials ($M$ = 96.2%, $SD$ = 3.4%, $t(15) = 26.54$, $p < .001$, $d$ = 6.64), and perceptual CoM occurred more frequently in test ($M$ = 10.3%, $SD$ = 9.9%) than easy trials ($M$ = 3.3%, $SD$ = 4.1%, $t(15) = 3.15$, $p = .007$, $d = 0.79$). Furthermore, in trials where participants did not have to verbalise their colour choice, accuracy showed a trend towards being lower in conflict ($M$ = 93.1%, $SD$ = 5.7%) than easy trials ($M$ = 95.7%, $SD$ = 3.5%; $t(15) = 1.89$, $p = .078$, $d = 0.47$). In contrast to Experiment 1, no difference in RTs was observed between conflict trials ($M$ = 560.8 ms, $SD$ = 47.5 ms) and easy trials ($M$ = 563.0 ms, $SD$ = 48.5 ms, $t(15) = 0.38$, $p = .712$, $d = 0.09$). However, the rate of misses (i.e., trials in which movement initiation exceeded the response deadline) was numerically increased in conflict trials ($M$ = 3.3%, $SD$ = 3.3%; easy trials: $M$ = 2.0%, $SD$ = 3.1%; $t(15) = 1.29$, $p = .216$, $d = 0.32$). Additionally, there were significantly more corrective movements in conflict trials ($M$ = 5.5%, $SD$ = 6.3%) than there were CoM in easy trials ($M$ = 3.4%, $SD$ = 5.5%, $t(15) = 3.51$, $p = .003$, $d = 0.88$) suggesting that, as in Experiment 1, participants generated initial colour intentions that resulted in response costs when external information did not match the endogenous intention.

#### S3 Attractor Network Model

**Table S3.1.** Fitted model parameters.

| Model parameters | Initial value | Fine-tuned value |
|---|---|---|
| *Connectivity weights w* | | |
| *w* intention → action | 1.00 | 0.97 |
| *w* sensory → action | 1.50 | 1.50 |
| *w* cost → action | -1.00 | -0.97 |
| *w* sensory → sensory | 0.25 | 0.25 |
| *w* lateral inhibition | -0.50 | -0.52 |
| *Decision evidence* | | |
| sensory evidence (% *coh*) | 3.20% | 1.03% |
| intentional strength (% *col*) | 51.2% | 48.0% |
| *Hierarchical noise control* | | |
| hierarchical control (*h*) | 1.00 | 2.01 |

**Table S3.2.** Model predictions and participants' actual behaviour in Exp. 1 [*M* (*SD*)].

| Outcome variable | Model prediction | Behaviour (Exp. 1) |
|---|---|---|
| Reaction times (ms) | 645.7 (4.9) | 570.5 (58.3) |
| % CoMov | 6.33 (0.8) | 5.93 (5.5) |
| % CoMov+Int | 1.41 (0.4) | 1.71 (2.2) |
| % Vertical color change | 3.58 (0.6) | 3.24 (2.6) |
| % double CoM | 1.61 (0.4) | 0.83 (1.0) |
| % perceptual choice accuracy | 54.5 (1.5) | 56.6 (9.1) |
| % color choice accuracy | 95.6 (0.6) | – |
| % misses (RT > 1000 ms) | 8.6 (0.9) | 9.3 (6.5) |
| % early responses (before stimulus onset) | 0.4 (0.2) | – |

### S4 Single-trial simulation

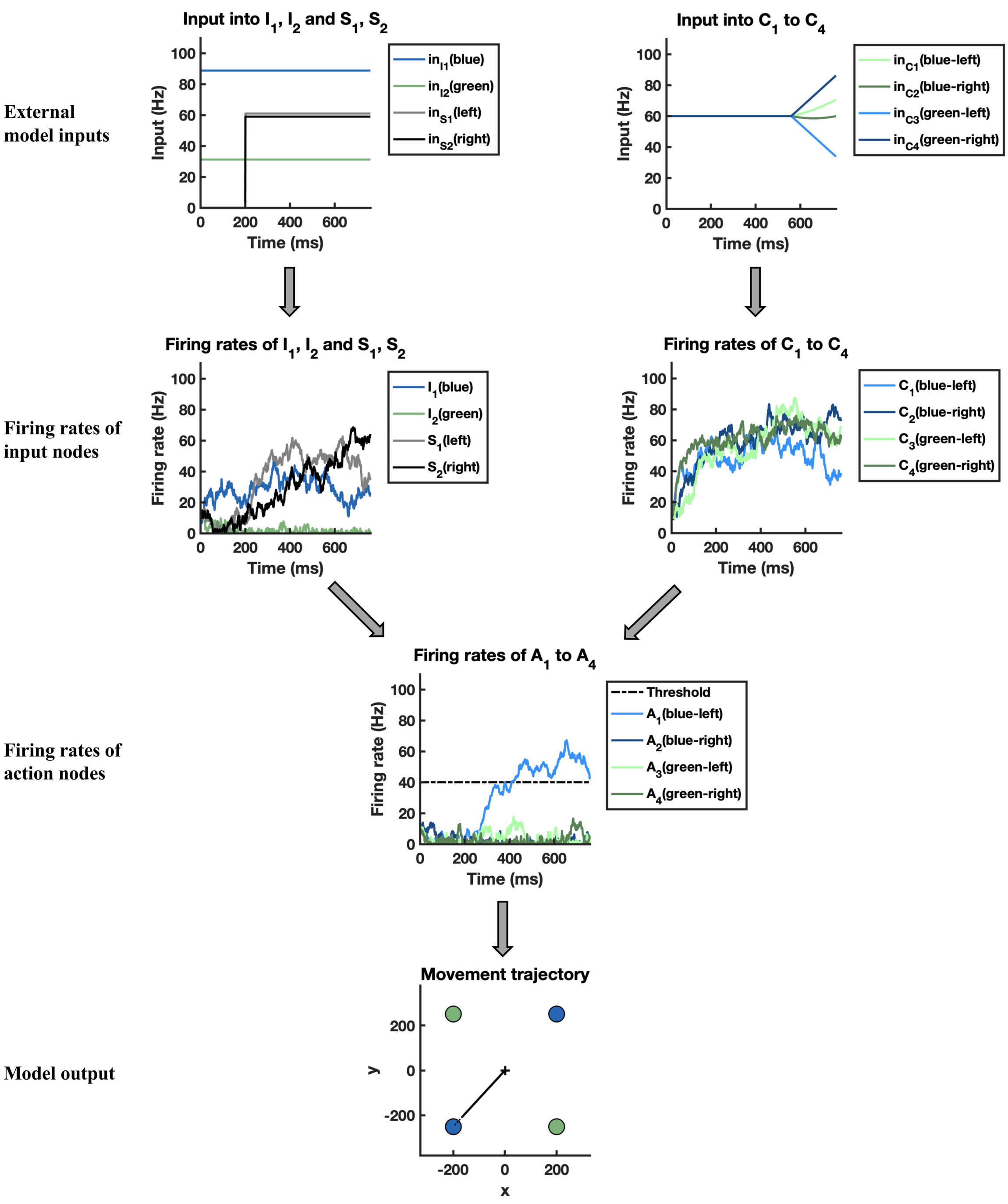


**Figure S4.1.** Simulation of a single trial without CoM. External model inputs (top row) are applied to intention, sensory and cost nodes (second row). Information is then combined by action nodes (third row) that determine the resulting movement trajectory (bottom row). In this example, the model correctly selects the left-blue target. After threshold crossing, the corresponding movement is initiated with a motor delay of 180 ms, and firing rates continue to be updated for a total non-decision time of 380 ms. Furthermore, inputs into cost nodes change after action initiation, according to the distance of the current cursor position to each target. Note that updates in cost nodes lag behind updates in action nodes due to the motor delay (i.e., costs only start changing 180 ms after a given action node has crossed the threshold).

### S5 CoM in ANM

##### *S5.1 Simulating the effect of intentional strength on CoM*

In Exp. 1 we found that participants with stronger colour intentions showed fewer CoMov+Int than participants with weaker colour intentions. To check whether the model could reproduce this finding, simulations with different degrees of intentional strength were performed (*col* = 0–65%), while all other model parameters were kept constant. In line with our behavioural findings, stronger colour intentions in the model predicted a decrease in the relative frequency of CoMov+Int out of all perceptual CoM (**Fig. 6B** in main text). Note that stronger colour intentions also caused fewer initial colour errors (**Fig. S6.1A**). However, even when trials with colour errors were excluded, CoMov+Int decreased relative to CoMov (**Fig. S6.1C**). This was due to the fact that stronger colour intentions shifted the percentage of colour errors that were corrected disproportionally towards vertical movement corrections (between targets of different colour on the *same* side of the screen), whereas CoMov+Int that corrected initial colour errors were less frequent for stronger intentions (**Fig. S6.1B**). Hence, the effect of intentional strength on CoMov+Int was not mediated by differences in initial colour errors.

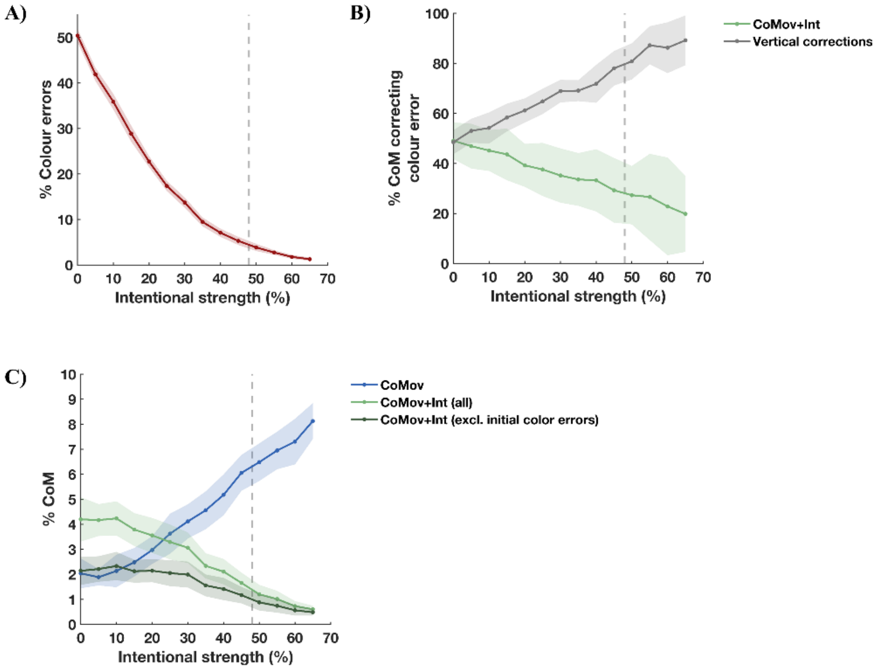


**Figure S6.1.** Effect of intentional strength on CoM in attractor network. A) Stronger colour intentions reduce % initial colour errors. B) Stronger colour intentions reduce % CoMov+Int (green), even when trials with initial colour errors are excluded (dark green). C) Stronger colour intentions result in more colour errors being corrected with vertical movement corrections as opposed to CoMov+Int [$M \pm 1$ *SD*].

##### *S5.1.1 Simulating conflict trials*

To simulate the effect of intentional strength on RT costs in conflict trials (**Fig. 6B** in main text), conflict and easy trials were simulated. For conflict trials, the model was changed such that each intention node mapped onto the two targets on the same (left/right) side (i.e., blue → both left actions $A_1$ and $A_3$; green → both right actions $A_2$ and $A_4$). The true colour intention of a given trial was selected randomly to induce a mismatch between colour intention and perceptual input on ~50% of trials (e.g., intention = green and both green targets are on the right side, but dot-motion direction = left). Additionally, in line with the behavioural task, both conflict and easy trials were simulated with a high motion coherence level (*coh* = 50%) to ensure that differences in RTs are not driven by perceptual difficulty. Note however that the precise level of motion coherence is of little importance for this simulation since RT costs are relative, i.e., they reflect the difference in RTs between conflict and easy trials (for correct perceptual decisions).

##### *S5.2 Simulating the effect of target distance on CoM*

The effect of target distance was simulated by changing the ratio of x:y target coordinates, while keeping all other model parameters constant. For example, in Exp. 2, target distances from the centre were $x = 300$ and $y = 100$, indicating a ratio of 3:1, and thus a far horizontal distance that causes low *relative* costs of intention pursuit. By contrast, a ratio of 1:3 ($x = 100$ and $y = 300$) indicates close horizontal targets, and thus a high relative cost of intention pursuit that should increase the frequency of CoMov+Int. Simulations with ratios ranging from 4:1 to 1:4 were performed. Crucially, the overall distance from the centre was equal for all targets, and hence, initial costs of each action were constant across all simulations. However, the relative change in costs after action onset varied as a function of x:y ratios in target locations. In line with our behavioural results, the model predicted an increase in CoMov+Int with closer horizontal targets (**Fig. 6D** in main text). Importantly, target distance did not affect the rate of colour errors (**Fig. S6.2A**), and hence, excluding trials with initial colour errors yielded the same pattern of results (**Fig. S6.2B**). Thus, differences in CoMov+Int across the different target distance conditions were not driven by potential differences in correcting initial colour errors.

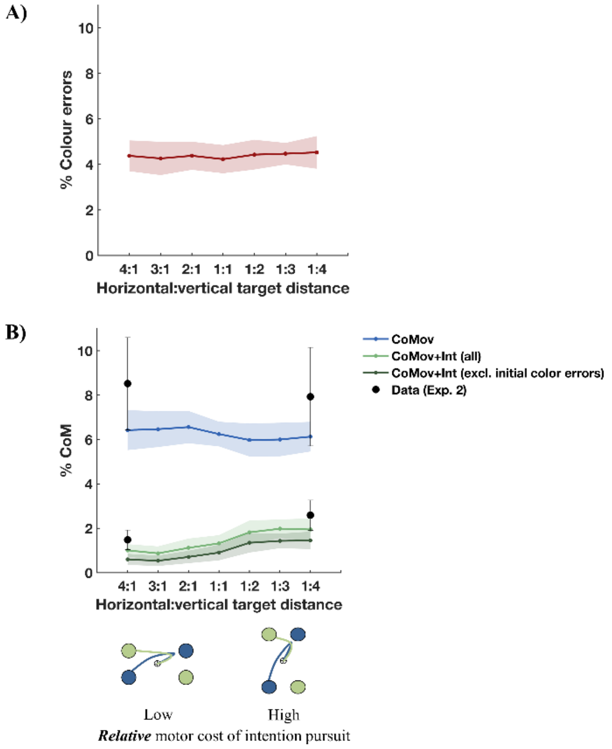


**Figure S6.2.** Effect of target distance (i.e., motor costs) on CoM in attractor network. A) Relative target distance does not affect % colour errors. B) Closer horizontal targets increase % CoMov+Int (green), even when trials with initial colour errors were excluded (dark green). Black data points represent behavioural results from Exp. 2 [$M \pm 1$ *SD*].
